# The comparative analysis of lineage-pair traits

**DOI:** 10.1101/2024.11.28.625927

**Authors:** Sean A. S. Anderson, Sachin Kaushik, Daniel R. Matute

## Abstract

A powerful but poorly understood analysis in ecology and evolutionary biology is the comparative study of lineage-pair traits. “Lineage-pair traits” are characters like ‘diet niche overlap’ and ‘strength of reproductive isolation’ that are defined for pairs of lineages instead of individual taxa. Comparative tests for causal relationships among such variables have led to groundbreaking insights in several classic studies, but the statistical validity of these analyses has been unclear due to the complex dependency structure of the data. Specifically, lineage-pair datasets contain non-independent observations, but studies to-date have relied on untested workarounds for data dependency rather than direct models of linear-pair covariance, and the statistical consequences of non-independence have not been thoroughly explored. Here we consider how evolutionary relatedness among taxa translates into non-independence among taxonomic pairs. We develop models by which phylogenetic signal in an underlying character generates covariance among pairs in a lineage-pair trait. We incorporate the resulting lineage-pair covariance matrix into a modified version of phylogenetic generalized least squares and a new beta regression model suitable for bounded response variables. Both models outperform previous approaches in simulation tests. We re-analyze two empirical datasets and find dramatic improvements in model fit and, in the case of avian hybridization data, an even stronger relationship between pair age and RI than revealed by standard linear regression. We present a new tool, the R package *phylopairs*, to allow empiricists from a variety of biological fields to test relationships among pairwise-defined variables in a manner that is statistical robust and more straightforward to implement.

## Introduction

An indispensable tool in the modern study of ecology and evolutionary biology is the broad-scale comparative analysis – a class of investigation in which data collected from numerous lineages are leveraged to test general support for competing hypotheses. The great power of comparative studies is that they identify statistical signatures that transcend the idiosyncrasies of any particular study system; comparative analyses thus complement the fine-scale study of individual lineages to expand our scope of inference. But comparative data also create statistical challenges. Most notably, whereas common parametric statistics assume independence among observations or model residuals, data such as the mean trait values of different species are expected to covary due to shared evolutionary history (Felsenstein 1985, Revell 2010). Models that explain this covariance (and methods to account for it) were developed throughout the 1980’s and 90’s and have been continually refined and elaborated in the modern study of phylogenetic comparative methods (PCMs) (Felsenstein 1985, Grafen 1989, Harvey and Pagel 1991, Lynch 1991, Martins and Hansen 1997). These advancements have given today’s biologists a strong understanding of the statistical consequences of shared evolutionary history among data points, but one highly influential form of comparative study has yet to receive sustained attention from modelers.

Here we are referring to comparative studies in which the traits of interest are attributes of lineage pairs, not of individual taxa. Examples of such pairwise-defined metrics include the strength of reproductive isolation (RI), relative hybrid fitness, range overlap, the extent of introgression, and ecological metrics like competition coefficient or diet niche similarity. The comparative analysis of lineage-pair traits involves calculating these variables among pairwise combinations of a set of taxa (Fig. 1), which distinguishes this approach from the study of non-overlapping sister pairs. Researchers then test for causal relationships among two or more of these pairwise-defined traits (e.g. range overlap versus divergence time, Fitzpatrick and Turelli 2006; hybrid fertility versus midpoint latitude, Yukilevich 2013; strength of RI versus ecological distinctiveness, Funk et al. 2006).

**Fig 1.**
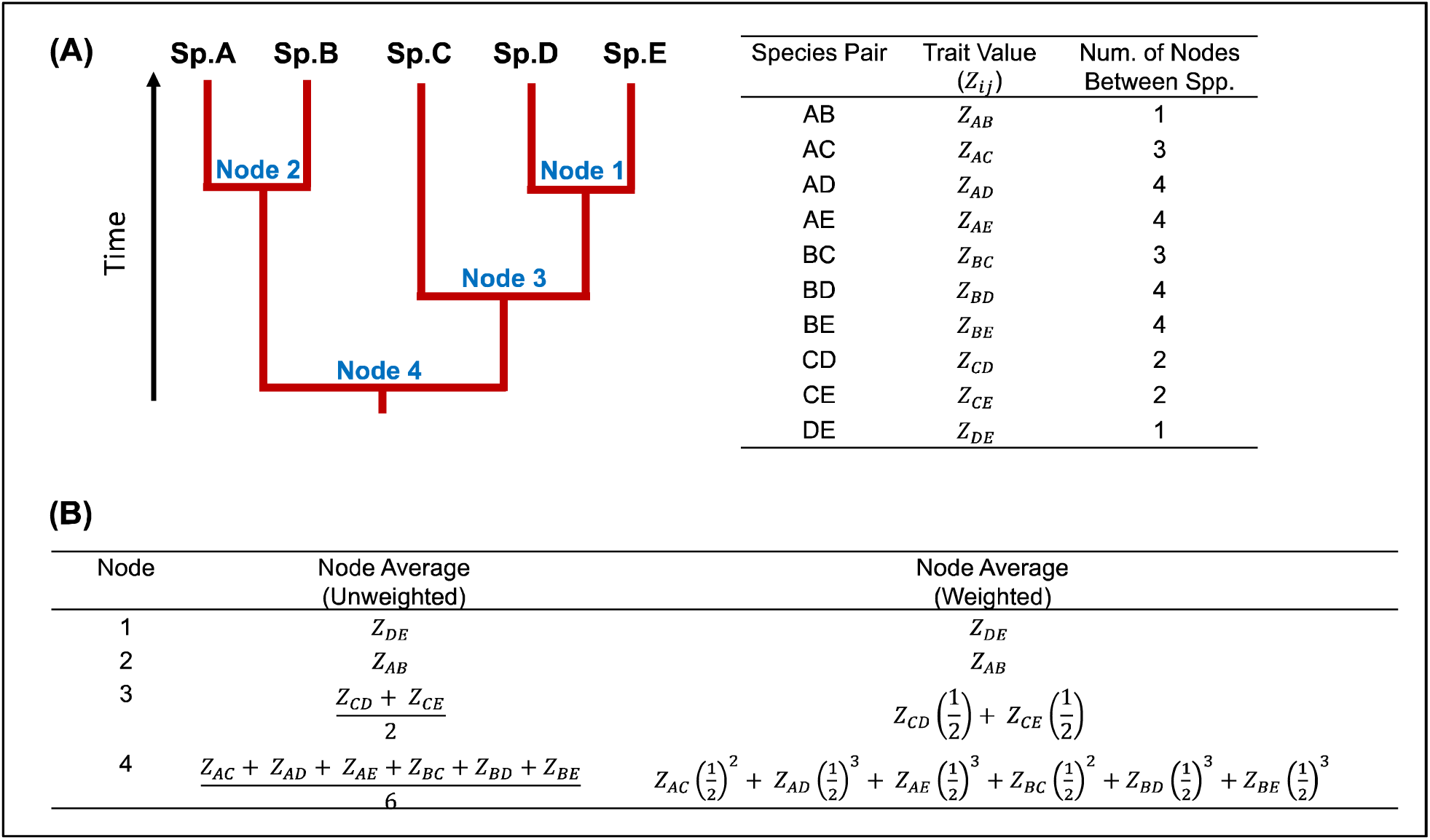
Deriving lineage pairs from a phylogeny and two methods of node averaging. A) a hypothetical phylogeny of five species. B) table showing all pairs that can be derived from species A-E in the phylogeny (i.e. all possible unique pairwise combinations of those taxa), where Z is the value of a hypothetical pairwise-defined trait for each pair. C) calculations of the unweighted and weighted average values of trait Z at each node of the tree. In the weighted average calculation, the exponent to which the (½) is raised in each term is K-1, where K is the number of nodes between species. Note that the number of data points obtained from node averaging (4) is less than half the number of actual trait measurements in the dataset (10).

To date, the analysis of lineage-pair data has represented a small but impactful subset of comparative biology. Its influence has been especially acute in the study of speciation, a field in which taxonomic pairs tend to make for natural units of analysis. Two pioneering works by Coyne and Orr (1989, 1997) on the rate of RI accumulation among more than 119 *Drosophila* species pairs are now classics in the field and have served as templates for today’s rich subgenre of comparative speciation research (Sasa et al. 1998, Presgraves 2002, Price and Bouvier 2002, Lijtmaer et al. 2003, Mendelson 2003, Russell 2003, Moyle et al. 2004, Bolnick and Near 2005, Malone and Fontenot 2008, Jewell et al. 2012, Owens and Riesberg 2013, Christie and Strauss 2018). Collectively, these comparative analyses of pairwise-defined traits have been enormously successful in helping to establish some of the very few general rules in evolutionary biology, including the modern axioms that the strength of RI increases monotonically with divergence time and that hybrid sterility tends to evolve prior to inviability (reviewed in Edmands 2002, Matute and Cooper 2021). But despite the outsized impact of lineage-pair analyses, existing studies have not benefited from a reliable and effective method for handling the underlying non-independence of their data. The consequences of this methodological limitation are not yet clear.

Like the traits of different species, the traits of different species pairs are not independent. Unlike the traits of species, we currently lack models that explain the structure of non-independence among species-pair traits. This makes it unclear how to properly account for their dependency. Indeed, most previous analyses of pairwise-defined traits have attempted to address non-independence in one way or another, but as we will outline in this paper, each previous effort at phylogenetic accounting suffers from one or more drawback including reduced statistical power and biased parameter estimation. It can also be practically challenging to implement these procedures, as standard computational tools for PCMs do not include options for working with pairwise-defined traits, thus custom computational scripts are typically required (though we note that ecologists have encoded methods to analyze spatial association in pairs of taxa from different guilds while incorporating, separately, the phylogenies for those groups, e.g. Pearse et al. 2015). Finally, and importantly, no previous approach has been subjected to a standard battery of model performance tests, which brings the statistical validity of some classic results into question. We suggest these drawbacks have limited the broader adoption of the comparative analysis of lineage-pair traits – though we maintain that there is great potential for the much wider application of this type of study if sound statistical approaches can be developed and if tools for their implementation are made available.

### The Problem: Pairwise-defined Traits Pose Unique Statistical Challenges

Three challenges arise in the comparative analysis of lineage-pair traits that differ from those of more conventional comparative analyses. The first is the unclear dependency structure of the data. In comparative studies of species’ traits, phylogenetic relatedness has a straightforward definition based on common ancestry, but there is no similar definition that dictates if one set of lineage pairs is more closely ‘related’ to each other than another. There are instead numerous conceivable ways to combine and weight the phylogenetic relationships among the (up to) four species in each pair of pairs, and each approach implies a model of evolution that must be clearly stated. Lineage-pair traits further introduce a second source of non-independence via the appearance of the same taxa in multiple pairs (Fig 1A.).

A second challenge posed by lineage-pair analyses is the bounded nature of many pairwise-defined traits. Metrics like the strength of (RI), range overlap, and competition coefficient are continuous but are typically bounded, often between 0 and 1. Basic linear regression procedures like Ordinary Least Squares (OLS) are therefore unsuitable for analyzing such data, as boundedness in a response variable violates the assumption of homoscedasticity (there will always be comparatively little variation around model-predicted trait values that lie near the bounds). Any clustering of observations near or on these bounds further complicates the calculation of least squares, which makes model fitting less reliable and can bias estimated regression coefficients.

A third challenge to lineage-pair analyses is that of “missing” data. By “missing” we are referring to pairs that can theoretically be formed from the species included in an analysis but that are not included in a lineage-pair dataset. If the value of a pairwise-defined trait is commutative (i.e. the trait for pair A-B is the same for B-A), then the number of possible pairs among N taxa is equivalent to the number of unique pairwise combinations of N (i.e. 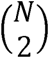). This number exceeds N in all datasets with four or more taxa and by rapidly increasing amounts as N gets larger. For analyses containing numerous species, it will be practically infeasible if not physically impossible to obtain data for each potential pair. For example, Price and Bouvier (2002) compiled all available data on avian hybrid crosses to amass a dataset of 407 crosses involving 368 species. This dataset remains one of the largest ever compiled for hybridization in animals, and yet its 407 crosses represent less than 1% of the 67,528 possible pairs that can be made from 368 taxa. Clearly, obtaining anything close to an additional 67,000 experimental crosses among vertebrate species is not practicable.

Effective methods to account for non-independence in lineage-pair datasets must therefore accomplish three things: (1) they must clearly define a model of lineage-pair covariance that also takes into account the appearance of the same taxa in multiple pairs, (2) they must accommodate bounded response variables, and (3) they must introduce no bias when fitted to datasets with large proportions of missing data. As we will demonstrate in the following sections, these requirements have not been met by the methodological approaches used to-date in comparative analyses of lineage-pair data.

### Previous Approaches to Account for Non-independence in Species-Pair Traits: A Review

Given the unique and unclear dependency structure of pairwise-defined data, the comparative analysis of lineage-pair traits cannot be conducted using the standard and powerful PCMs that have been devised for the comparative study of species’ traits. Researchers have therefore employed one of two custom approaches to account for non-independence in lineage-pair datasets: node averaging and a modified version of a phylogenetic linear mixed model.

#### Node Averaging

The most common approach to account for non-independence in species-pair datasets has been the method of node averaging. Coyne and Orr (1989) first introduced an unweighted node averaging procedure in which they calculated, at each node in the tree, the average value of a pairwise-defined trait for all pairs whose species spanned the node (Fig. 1). The logic of node averaging is that the resulting mean value at a node is independent of the mean values at other nodes, a concept borrowed from Phylogenetic Independent Contrasts (PICs, Felsenstein 1985). Node averaging has consequently been referred to as a special form of PIC in subsequent papers (e.g., Fitzpatrick 2002, Turrelli et al. 2014, Castillo 2017), but this is a misnomer: node averaging is mathematically unrelated to Felsenstein’s independent contrasts and is in fact not a contrast at all (i.e., no subtraction occurs). A more substantial difference is that PICs follow from a formal theoretical basis whereas node averaging does not. In PICs, a continuous trait is modeled as evolving via Brownian motion (BM) independently in each branch of a phylogenetic tree. The difference, or ‘contrast’, in the traits of two lineages is then the difference in the outcome of two independent random walks. This difference in a normal random variable whose variance depends only on the BM scaling parameter and the time since a common ancestor. The distribution of each contrast is thus independent of all other contrasts. Node averaging also produces values that are independent, but the averaging itself is heuristic and has no similar theoretical underpinning.

Node-averaging, while useful and widely adopted, presents three drawbacks. First, node averaging does not take phylogenetic branch lengths into consideration. A central tenet of modern PCMs is that the expected covariance between two taxa depends on the amount of evolutionary history they have shared, and this history is represented by the total length of their shared branches in a phylogenetic tree. By ignoring branch lengths, node-averaging does not account for varying amounts of shared evolutionary history among lineages and so does not achieve the principal goal of most phylogenetic corrections. Second, unweighted node-averaging overlooks some proportion of total non-independence among pairs. For example, when calculating the value for Node 4 in Fig. 1A, we treat pairs AC, AD, and AE with equivalent weights and average their values. But while species A is equally related to species C, D, and E, we can also see that species D and E are more closely related to each other than either is to species C. We might then expect AD and AE to have more similar pairwise-defined trait values, but this information is lost in the averaging.

A third problem with node averaging is statistical power loss. A fully bifurcating tree with *N* species contains *N*−1 nodes. We previously noted that the number of possible pairs among *N* species is 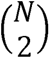, which greatly exceeds *N* in most datasets. Much statistical power is therefore lost by collapsing a dataset of size 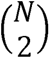 down to the *N*−1 nodes in a tree. In practice, few datasets contain data for all 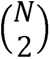 possible pairs, but the problem of power loss remains nonetheless: node averaging in Coyne and Orr (1997), for example, reduced a dataset of 171 species pairs down to 50 node-mean values of postzygotic isolation. Given the substantial power loss induced by node averaging, studies employing this approach typically report results from both “uncorrected” and node-averaged data. While this practice of dual reporting was once routine in phylogenetically informed analyses, it is no longer considered the best practice due to the fundamentally incompatible assumptions that the different analyses make about data structure (Pagel 1999, Revell 2010, Symonds and Blomberg 2014).

An improvement upon unweighted node-averaging was introduced by Fitzpatrick (2002), whose weighted node-averaging procedure takes into account phylogenetic nesting. In Fitzpatrick’s (2002) approach, the trait values of the pairs spanning a given node are each halved *K*-1 times, where *K* is the total number of nodes between the taxa in a given pair. These reduced values are then summed to produce a weighted average (Fig 1.B and see Fig. 1 of Fitzpatrick and Turelli 2006). This procedure has the desirable effect of down-weighting certain combinations of pairs whose respective taxa are closely related. For example, when calculating the value for node 4 in Fig. 1B, the traits for AC, AD, and AE are no longer simply averaged; AD and AE are instead down-weighted relative to AC due to the additional node between A and E and between A and D. But despite its improvement over the unweighted approach, weighted node averaging still results in substantial power loss and does not incorporate branch lengths. Moreover, as we will find in the section on Model Performance, this method can introduce a significant bias when datasets are missing observations for at least some possible pairs.

#### The Modified Phylogenetic Linear Mixed Model

A more recent approach to analyzing lineage-pair data is a modified version of the phylogenetic linear mixed model (PLMM; Castillo 2017). A standard PLMM is a type of linear mixed model (LMM) in which the random effects covary according to the structure of a phylogenetic covariance matrix. Castillo (2017) modified the basic version of the PLMM to make it more suitable for lineage-pair data by defining two random-effect terms, one for each species in every pair. The trait values of different pairs are thus expected to cluster based on the identity of their “Species 1” and “Species 2”, where the distinction between the two may or may not have biological significance (e.g., in hybridization studies, Species 1 may be the female species and Species 2 the male species of each pair; in other studies, the two sets may be arbitrarily defined). We refer to this model hereafter as the two-term LMM. Castillo’s (2017) approach offers several substantial advantages over node averaging; namely, no power is lost, the approach can be readily adapted for models with multiple predictor variables, the models can be implemented by slight modifications to existing computational tools (e.g. MCMCglmm, Hadfield 2010), and phylogenetic branch lengths are brought into consideration via the covariance matrix for the random-effect terms (but note that a pairwise genetic distance matrix is sometimes used in place of a phylogenetic covariance matrix, in which case branch lengths are not considered).

What is unclear about the two-term LMM is the underlying evolutionary model that it implies and how to interpret that model biologically. Specifically, under what model of evolution might we expect to see the two-term LMM’s unique clustering structure in a lineage-pair trait? We note that some amount of phylogenetic covariance among pairs gets compressed by the act of separately modeling the random effects of Species 1s and Species 2s, and it is unclear if this should be expected biologically. Among the pairs that can be generated from the tree in Fig. 3, for example, are AC, AD, BC, and BD. The pairs AC and AD share a “Species 1” but have a different “Species 2”. The covariance in their Species 1 random effects is then proportional to the branch lengths of the species tree from the root to the tip A, while the covariance in their Species 2 random effects is based on the shared branch lengths of species C and D. This exact situation is also true for the pairs BC and BD, and since species A and B are both extant, the two pairs will have identical covariances in both the Species 1 and Species 2 random effects (Fig. 3C). That is, the combined covariance among random effects between AC and AD is identical to that for BC and BD, despite the fact that B is more closely related to C and D than is A.

**Fig. 2.**
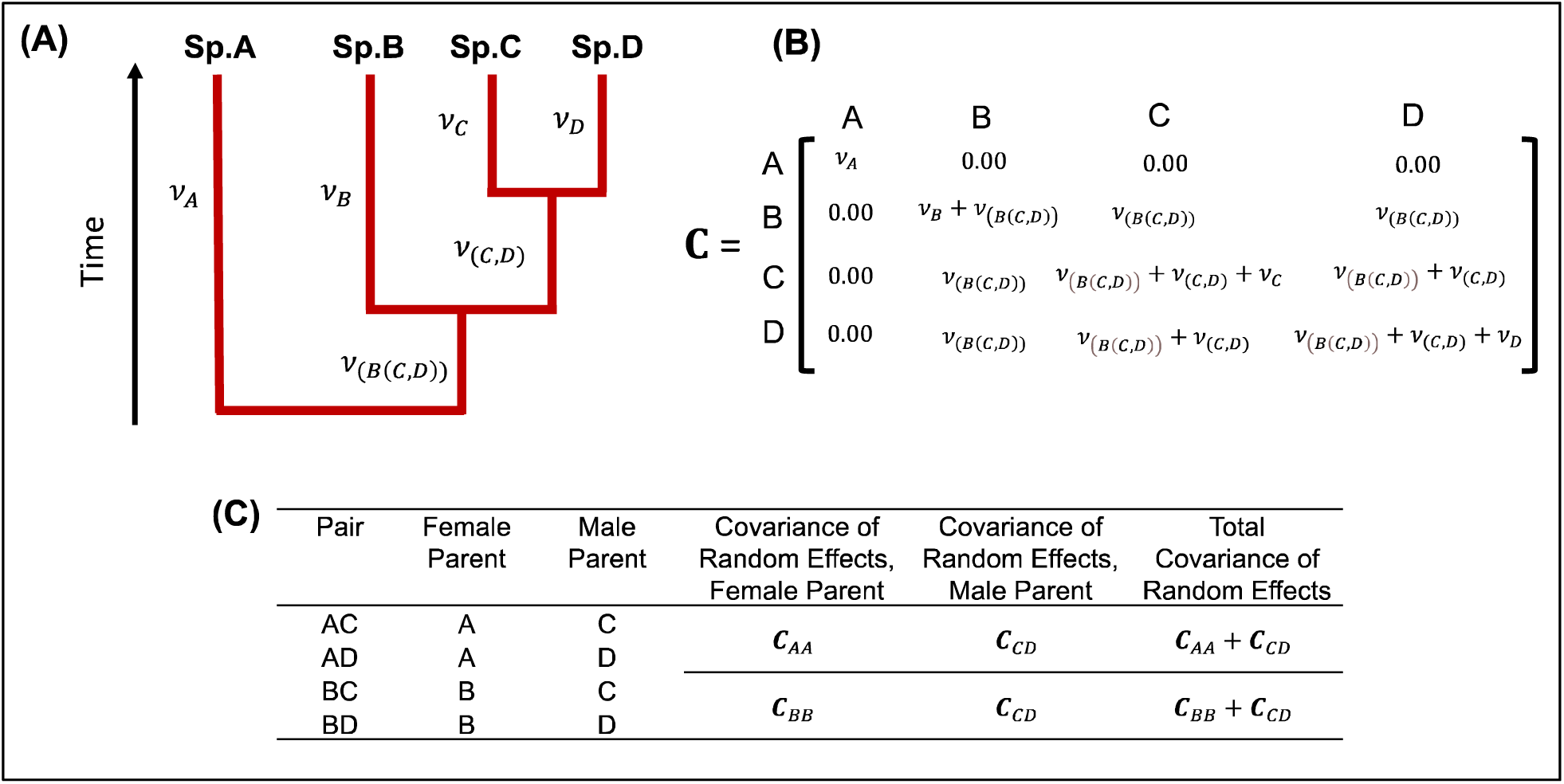
Consequences of separating random effects of “Species 1” and “Species 2” in each pair. A) a phylogeny of four species with branch lengths. B) the phylogenetic covariance matrix **C**, which defines covariance of both the “Species 1” and “Species 2” random effects in Castillo’s (2017) two-term LMM. C) calculating the covariance of random effects for two sets of pairs, AC-AD and BC-BD, when maternal and paternal effects are considered separately in a two=term LMM. **C**_AA_ and **C**_BB_ are equivalent when species A and B are extant, which means that the total covariance in random effects are identical among the two sets of pairs despite the difference in their overall phylogenetic relatedness: species B is more closely related to C and D than is species A.

**Fig 3.**
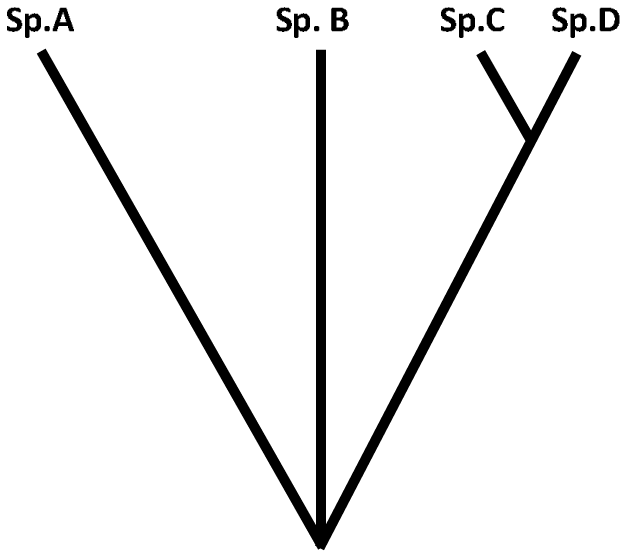
A four-species tree. Six pairs can be derived from the taxa in this tree, including AB, AC, and AD. In the main text, we consider whether we would expect any two of these three pairs to be more similar to each other than either is to the third in some pairwise-defined trait.

### Translating Phylogenetic Signal into Lineage-Pair Covariance

A fundamental shortcoming of the existing methods for dealing with non-independence in lineage-pair data is that none attempts to calculate the expected covariance among the pairs themselves. To test for evolutionary relationships among pairwise-defined traits in a manner that properly accounts for lineage-pair non-independence, we must first clarify how the traits of different lineage pairs are expected to covary.

To this end, it helps to interrogate why we expect phylogenetic relationships to matter in a lineage-pair analysis. Consider the tree in Fig. 4. Of the four species A, B, C, and D, we can derive six pairs, three of which are AB, AC, and AD. A useful question to consider is if we should expect any two of these three pairs to be more similar to each other in some continuous pairwise-defined trait than either is to the third.

**Figure 4.**
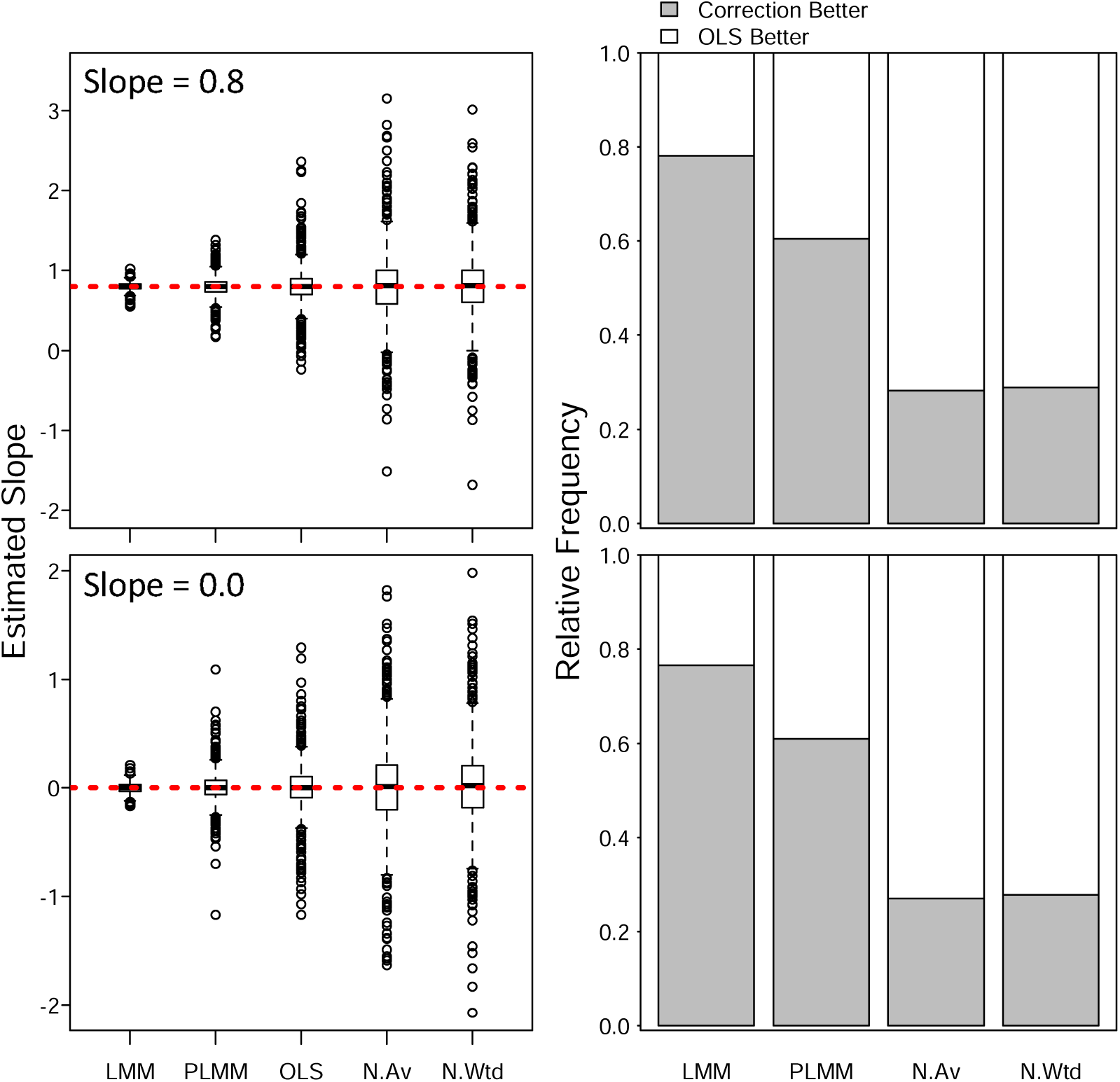
Estimating regression coefficients in lineage-pair data. Values in the left column are replicate means of posterior distributions for Bayesian-estimated regression coefficients. Dashed red lines show the regression slopes with which datasets were simulated; slope values apply to the entire row. Each model was fit to 1000 simulated datasets with unbounded responses. All models were given the same prior distributions of *b*_1_∼*N*(0,10). “N.Wtd” and “N.Av” indicate weighted and unweighted node averaging, respectively. The lineage-pair LMM, the two-term LMM, and OLS models were fit to datasets with 190 observations. Node averaging condenses these datasets to the number of nodes in the tree (19). Y-axes in the left column are not equivalent. Stacked bar graphs in the right column show the efficacy of four different corrections for data non-independence. Values are proportions of replicate analyses in which a standard OLS regression or the given correction returned a mean slope estimate closer to the true value.

An intuitive expectation might be that AC and AD are more similar to each other in a pairwise-defined trait than either is to AB. For example, if species A has strong postzygotic RI with species C, then we might expect that it also has fairly strong postzygotic RI with species D. A biological explanation might be that since postzygotic RI arises from incompatibilities in the genomes of two species, then species A will tend to be similarity compatible (or incompatible) with species C and D, as the latter have only recently diverged and will likely have fairly similar genomes. Similarly, if we are analyzing an ecological metric like competition coefficient, we might intuit that if species A competes strongly for resources with species C, then it also competes fairly strongly for resources with species D.

Our basis for these expectations is the implicit assumption that C and D are themselves similar in some trait or traits due to their shared evolutionary history, and this similarity affects the value of pairwise-defined traits of both species with species A. In other words, we are expecting *phylogenetic signal* in some underlying character or set of characters that influences the pairwise-defined trait we wish to analyze. The operational challenge is to translate the phylogenetic signal in the underlying influencer trait(s) into the expected covariance among pairs in our pairwise-defined character of interest, and to do so without having to know what those underlying characters are.

### Model Description 1: Trait “X” and the Lineage Pair Covariance Matrix **C_P_**

Following our intuition from Fig. 3, we state that the value of a pairwise-defined trait is influenced by an underlying and unmeasured biological character that has phylogenetic signal. We model this underlying character as a continuous trait, *X*, which evolves through the branches of a species tree under BM (the expectation of phylogenetic signal). The value of *X* in any lineage is then a normal random variable, and the covariance among species in *X* is described by a phylogenetic variance-covariance matrix, **C**, scaled by the BM dispersion parameter 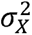. We can think of *X* as being any biological character or composite of characters that tends to be similar in closely related species and that somehow influences the species-pair trait that we care about. How that influence is exerted will determine the expected covariance structure in the pairwise-defined trait.

We first consider a simple model in which the lineage-pair trait is affected by the magnitude of the difference between species in *X* (e.g. if the trait is postzygotic RI, then *X* may be some aspect of genomic content, and we expect the value of RI to increase monotonically with the magnitude of the difference in *X*; if the pairwise-defined trait is something like competition coefficient, then *X* may be the value on some axis of the ecological niche, and we expect the value of the lineage-pair trait to decrease monotonically with the magnitude of the difference in *X*). This magnitude is itself a random variable *Y*, calculated as the square difference between species in *X*, where 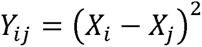.

For two lineage pairs, one comprised of taxa *i* and *j* and the other of taxa *k* and *l*, the covariance between pairs in *Y* is given by

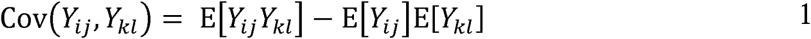

If we consider the ancestral value of *X* at the root of the tree to be zero (i.e. *X*_A_ = 0), then it follows from the assumption of BM evolution that *X_i_*, *X_j_*, *X_k_*, and *X_l_* are jointly distributed zero-mean normal random variables. By Isserlis’ theorem, we can then express the unscaled covariance among pairs in *Y* in terms of the unscaled covariances among species in the influencer trait *X* (Appendix 1). The covariance in Equation 1 then expands and simplifies to:

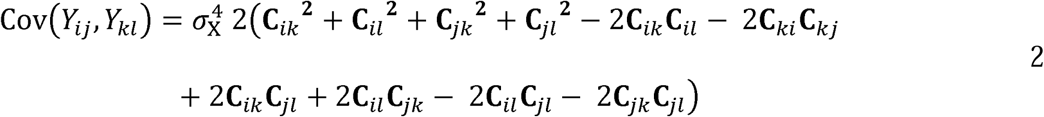

In the R package *phylopairs*, the function ‘taxapair.vcv’ calculates the unscaled version of this covariance (i.e. equation 2 not scaled by 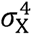) for each pair of pairs in a dataset to generate a lineage-pair variance-covariance matrix that we hereafter denote **C***_P_*. If there are N species in a dataset, then a standard phylogenetic covariance matrix **C** has dimensions N × N, and the lineage-pair covariance matrix **C***_P_* has dimensions 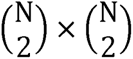. We remind readers that *Y* is not the lineage-pair trait we are analyzing; rather, the trait we are analyzing is some function of *Y*. We will assume that this function is linear, which makes the covariance among pairs in our trait of interest (or the covariance of the residuals when this trait is regressed against some set of predictor variables) equivalent to **C***_P_* scaled by a parameter that we estimate from the data.

We have assumed that the pairwise-defined trait is influenced by the magnitude of the difference between taxa in *X*, but there are conceivable biological scenarios in which that trait might instead be more strongly influenced by the sums or the products of *X* in each species (i.e. . 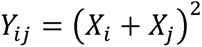 and 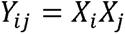). Such scenarios would result in different covariance structures, and which (if any) of these hypothesized covariances matches that of a given dataset is ultimately an empirical question. In the package *phylopairs*, users can calculate **C***_P_* based on the each of these three models and compare the fit of a given model to a dataset when assuming the alternative covariance structures. Formulae for the expected covariance resulting from each model are shown in Appendix A. For the remainder of this paper, all simulation-based tests will be conducted using the first model: 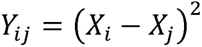.

### Model Description 2: The **C_*P*_** Matrix in Regression Analyses of Pairwise-Defined Traits

The lineage-pair covariance matrix, **C**_*P*_, can now be used to account for non-independence in any number of analyses based on pairwise-defined data. Here we introduce two regression-style approaches, one for unbounded and one for bounded responses.

#### The Lineage-Pair Generalized Least Squares (LPGLS) Model for Unbounded Reponses

Castillo’s (2017) framework was based on a modification of the LMM that had two terms for phylogenetically non-independent random-effects, one for each of the two species in every pair. He we consider a similar model that instead has a single random effects term. Specifically, each pair is modelled as having its own random intercept where the intercepts are not independent but covary according to **C**_*P*_. The model is:

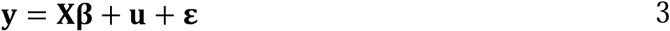

where ***y*** is an *m* × 1 response vector of *m* observations; **β** is a *p*×1 the vector of unknown regression coefficients 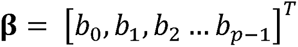, where *b*_0_ is the intercept and *b_x_* is the regression coefficient for the *x^th^* predictor); **X** is an *m* × *p* design matrix that maps *p – 1* predictors to their coefficients in **β** (the first column in X contains 1s and maps to the intercept); **u** is an *m* × 1 vector of pair-specific random effects where 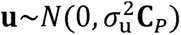; and **ε** is an *m* × 1 vector of model residuals where 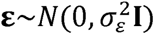.

The model in equation 3 is a form of generalized least squares (GLS) regression. In both OLS and GLS, the model is defined as **y = Xβ + ε**. In OLS, the residuals are assumed to be independent and identically distributed Gaussian variables with mean zero and constant variance 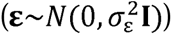. GLS relaxes those assumptions by allowing the residuals to have a known, invertible covariance matrix 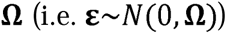. Our model is equivalent to a GLS in which 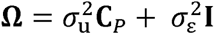. This model can also be considered an extension of Phylogenetic Generalized Least Squares (PGLS). PGLS is an adaptation of GLS for analyzing species’ traits in a phylogenetically informed context (Grafen 1989; Martins and Hansen 1997; Pagel 1997, 1999; Rohlf 2001; Revell 2010; Symonds and Blomberg 2014). PGLS is simply GLS in which **Ω** is the phylogenetic covariance matrix, **C**, scaled by a parameter 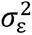. In equation 3, the random effects contain a lineage-pair version of that term in which 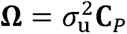 as well as uncorrelated residual error component. In the *phylopairs* package, users can fit the model in equation 3 as well as a simpler lineage-pair version of PGLS in which there is no additional residual error term.

#### Bounded Responses: A Mixed Model of Beta Regression for Phylogenetic Analyses of Bounded Data

A relatively new method for conducting regression analyses with bounded response variables (as will be the case in many comparative studies of lineage-pair traits) is beta regression (Ferrari and Cribari-Neto 2004, Cribari-Neto and Zeileis 2010). The two-parameter beta distribution for a variable bounded in the interval (0,1) can assume a wide variety of shapes within its bounds and is usually described by two positive “shape” parameters, typically denoted *α* and *β* or *p* and *q*, depending on the source. An alternative parameterization was created by Ferrari and Cribari-Neto (2004) to enable a regression analysis for bounded response variables. Instead of two positive shape parameters, the beta distribution is expressed by its mean, *µ*, and a “precision” parameter *φ*. The probability density function of this distribution is:

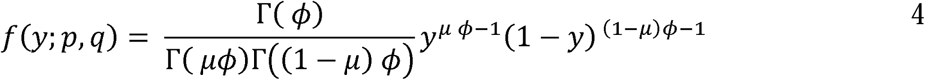

and we write 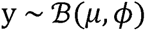 (Cribari-Neto and Zeileis 2010).

In its most basic form, beta regression assumes a linear relationship between response and predictors on the scale of a link function. If **X** is an *m × p* matrix of *m* observations of *p* − 1 predictors, and ***β*** is a *p ×* 1 vector of unknown regression coefficients, then the model is

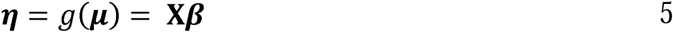

where ***η*** is an *m ×* 1 vector of predicted values of ***y*** on the scale of a strictly increasing link function *g*. Common choices of the link function include the logit and the probit, among several others (Cribari-Neto and Zeileis 2010). The predicted values of ***y*** from the model are then given by *g*^−1^(***η***).

We previously stated that a problem with using linear regression when analyzing bounded response variables is that the assumption of homoscedasticity is violated, as bounded variables will show less variation around the model’s predicted value as that value approaches the upper and lower bounds. Beta regression is naturally heteroskedastic and so avoids this problem. In beta regression, the variance about the predicted value is a function of the mean *µ*, where

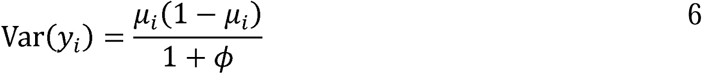

A key distinction between beta regression and more common forms of regression with continuous responses (including both linear and non-linear least-squares regression) is that residuals are not directly modeled. In OLS and GLS, the linear predictor includes a term for residual error (e.g. **ε** in **y = Xβ + ε**). The linear predictor in beta regression includes no such term. Instead, deviations from the predicted values are determined by *φ*, whose effect we can see in equation 7: larger values of *φ* result in less dispersion about the expected value of *y_i_*. This fundamental distinction changes how we incorporate the non-independence of lineage-pair data into the analysis. In linear regression, the key assumption being violated by phylogenetic relatedness is non-independence of the residuals. Beta regression does not share that assumption. However, beta regression does expect independence of observations, and violations of this assumption can bias estimates of regression coefficients.

Our approach is therefore to create a mixed effects version of the beta regression model. Similar to the lineage-pair LMM above, the linear predictor is now

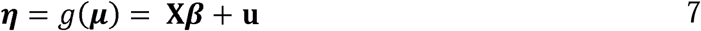

where **u** is an *m* × 1 vector of pair-specific random effects 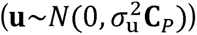

#### Fitting the Lineage-Pair LMM and Beta Mixed-Effects Model in ‘phylopairs’

The likelihood functions of the lineage-pair LMM and the beta mixed-effects model have no closed-form solutions, so we opted for a Bayesian approach to model fitting in all analyses to follow. Functions to fit both models are encoded in the *phylopairs* package. We wrote the Bayesian models as programs for the ‘Stan’ software program (Stan Development Team 2024) and call them within *phylopairs* via the *rstan* package (Stan Development Team 2024). Note that in a Bayesian context, there is no formal difference between fixed and random effects, but for clarity and consistency we will continue to use that terminology to distinguish between the impact of the predictor versus data non-independence on the response variable.

### Tests of Model Performance

To date, no method used in a comparative analysis of lineage-pair data has been validated in tests of model performance. The ability of these methods to accurately estimate regression coefficients and their sensitivity to missing data are therefore unknown. Here we conduct three series of model performance tests to determine both the consequences of non-independence in lineage-pair data on parameter estimation and the ability of our proposed approach to improve upon existing methods. Code for all performance tests is included as supplementary material. Package *phylopairs* is available for download from the Comprehensive R Archive Network.

#### Parameter Estimation: Unbounded Response

We began by assessing the ability of our linear mixed model (based on the lineage-pair covariance matrix, **C***_P_*) to improve the precision of parameter estimates in comparison with previous approaches. We will consider first an unbounded response variable so that the only assumption being violated by the lineage-pair relationships is that of independent residuals. Details on all simulation procedures are in the Materials and Methods. Results of parameter estimation are shown in Fig. 4.

The lineage-pair LMM performs best in terms of accuracy and precision in parameter estimation: the means of its posterior distributions cluster tightest around the true value (Fig. 4A,C) and its posterior distributions have the narrowest credible intervals (Fig. S1). Castillo’s (2017) two-term LMM also offers improvement over OLS, particularly when the true slope is nonzero, while the node averaging procedures generate more variable coefficient estimates. Only the lineage-pair LMM and the two-term LMM offer a consistent improvement over naive OLS regression (Fig. 4B,D). The results here suggests that node averaging should be avoided; the power loss induced by node averaging imparts a greater performance cost than does the statistical non-independence that the method is attempting to correct for.

#### Sensitivity to ‘Missing’ Data

An important test of any method used in lineage-pair analyses is sensitivity to data incompleteness. Here we revisit the five models tested above to compare the accuracy and precision of each model’s parameter estimations when fit to datasets with five increasing levels of missing data. By ‘level of missing data’ we are referring to the proportion of the total number of pairs that can be made from the species in a tree and that are not included in the final dataset. Results are shown in Fig 5.

**Figure 5.**
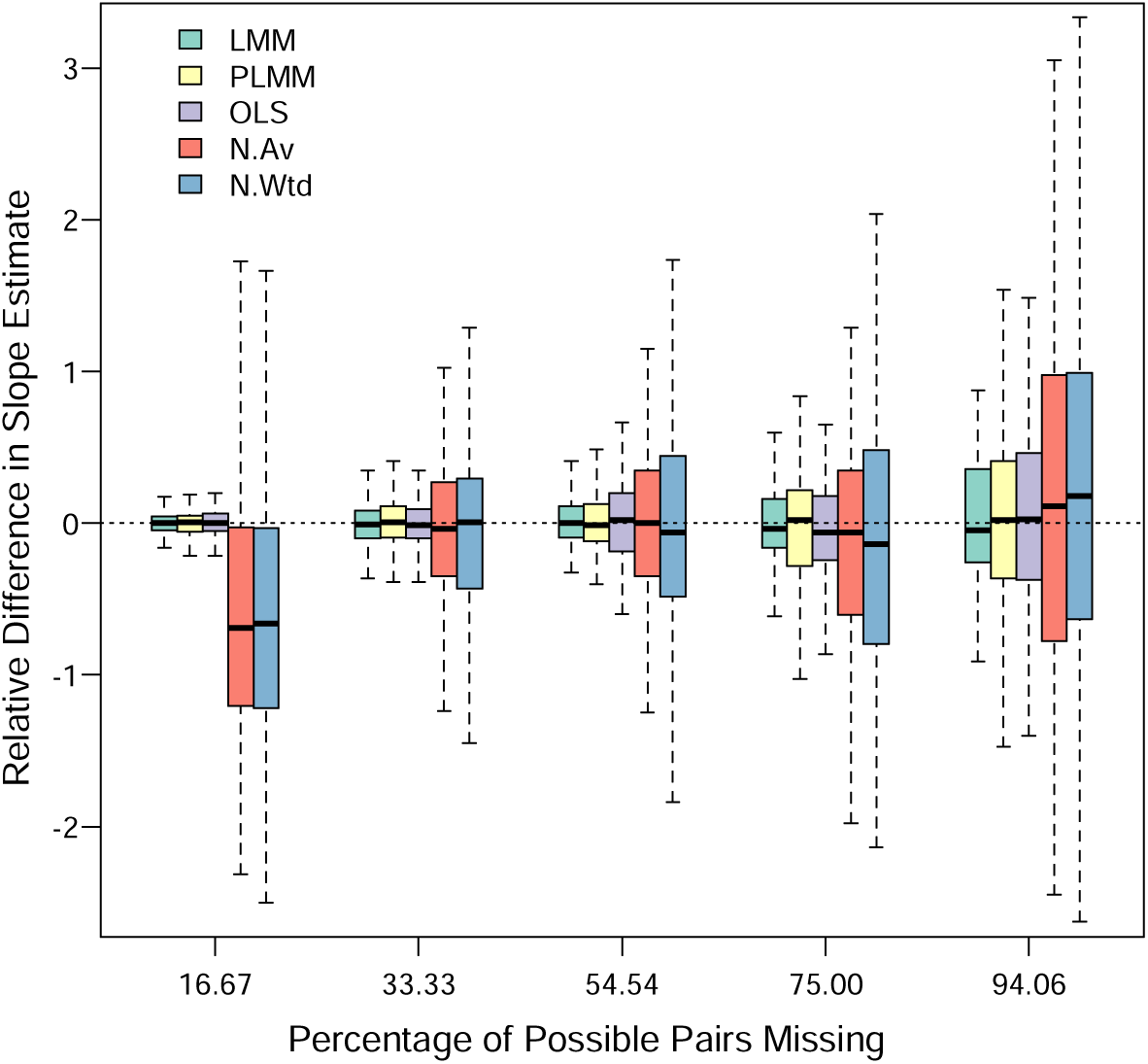
Effect of ‘missing’ data on slope estimation. Boxplots show the distributions of the differences in mean posterior regression coefficients estimated by each model when fit to incomplete versus complete datasets. Zero values correspond to no change in parameter estimate; positive and negative values that indicate higher and lower parameter values, respectively, were estimated from incomplete relative to complete datasets. Differences were scaled to be relative to the true slope (0.8). At each level of missing data, models were fit to 500 complete datasets and 500 datasets with some number of missing pairs, where that number represents the percentage shown on the X-axis. The size of the incomplete datasets were constant at 30 pairs across the five treatment levels for the lineage-pair LMM, the two-term LMM, and OLS models, while node averaging procedures (“N.Av” and “N.Wtd”) generated datasets of variable size when pairs were missing (see Supplementary Text).

For the lineage-pair LMM, Castillo’s (2017) two-term LMM, and OLS models, the relative variability in parameter estimation increased with increasing proportions of missing data, as expected (Fig. 5). The lineage-pair LMM was least affected by missing data overall. The two-term LMM was affected by missing data to an equal or greater extent as OLS at all levels of data missingness except Level 3. The central tendencies of the distributions of the three models in Fig. 5 vary little across levels and hover near zero, suggesting no systematic bias is induced by these models when fit to incomplete datasets. For the node averaging procedures, by contrast, relative variability in parameter estimation was greatest at data missingness Level 1, where a downward bias was also observed for both models. The variability and bias in estimates produced by node averaging procedures are likely the results of extremely low sample size. At Level 1 of data incompleteness, all trees contain 9 taxa and 8 internal nodes, thus node averaging reduces the sample size down from 30 to a maximum of 8. Any further decrease in that sample size due to missing pairs (see Supplementary Text) makes it unlikely that the model will have power to detect an existing slope. Weighted node averaging was affected to a slightly greater extent by missing data than the unweighted method at all treatment levels.

While not apparent in Fig. 5, we found that weighted node averaging creates a downward bias in the values calculated at each node when some possible pairs are missing in the dataset. This bias is masked in Fig. 5 because the predictor and response variables were affected in the same way, thus the regression coefficients were not systematically skewed. But the data vectors being regressed contained notably smaller values than those calculated from complete datasets.

We investigated this bias further by comparing the weighted node averages calculated from simulated datasets and from versions of those datasets in which varying levels of pairs were randomly removed (i.e. ‘missing’). In the results in Fig. 6A and 6B, we see that the weighted mean calculated for a node decreases relative to the true weighted mean when some proportion of pairs are not included in the analysis; this effect is more pronounced at deeper nodes in the tree and worsens with greater proportions of missing data. The systematic underestimate observed for weighted node averages is not observed for unweighted averages at any level of missing data.

**Figure 6.**
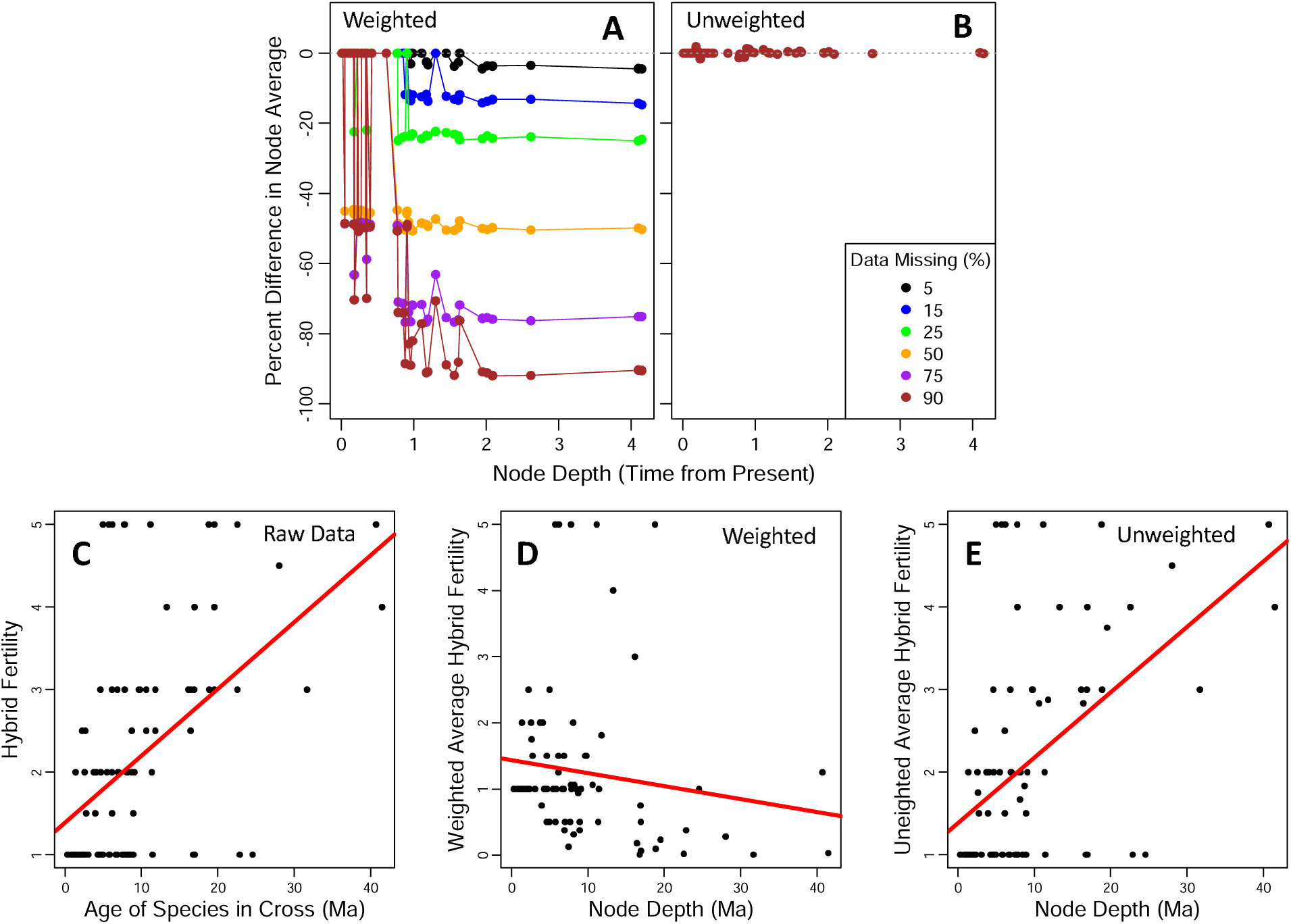
Bias in weighted node averaging. A-B) points are medians of 1,000 replicate errors in node-averaged values of an arbitrary trait calculated at each node in a tree of 50 taxa (49 nodes). ‘Error’ is the percent difference in the value calculated at given node from an incomplete dataset relative to the complete dataset. Weighted averages are shown at 6 levels of missing data; for clarity, we show unweighted averages only for the most severe missing data level. Note the systematic underestimate of weighted but not unweighted node averages at deeper nodes in the tree. C-D) an empirical relationship between the index of postzygotic RI and evolutionary time from avian hybrid crosses (Price and Bouvier 2002). Values are raw data (C), weighted (D) and unweighted node averages (E). Red lines are OLS regression estimates.

To understand the source of this error, recall that in the weighted node averaging procedure, the value of a trait for a pair whose species span a node is halved *K*−1 times, where *K* is the total number of nodes between the species (Fig. 1C). As we move to deeper and deeper nodes in the tree, we are averaging traits from pairs whose species are separated by an increasingly greater number of nodes. The trait values of these pairs are thus halved many more times relative to pairs that span shallower nodes in the tree. This poses no problem when the dataset is complete, because deeper nodes are also spanned by more pairs, and many of these small values add together to reconstitute a reasonable weighted average. But when numerous pairs are excluded from the analysis, we end up adding too few of these shrunken values together. The result is an erroneously small weighted mean calculated at the deeper nodes. This bias can have important implications for analyses in which only one variable is averaged. For instance, in many lineage-pair studies, divergence time is the independent variable. Divergence time is equivalent for all pairs spanning a node and is simply the node depth in a dated tree. Fig. 6C-D shows that when naively applied to Price and Bouvier’s (2002) hybridization dataset in this scenario, weighted node averaging of the response variable leads to the counterintuitive result that more anciently diverged species tend to generate more fertile hybrids. Presumably the weighted node averaging procedure can somehow be modified to counteract this trend.

#### Parameter Estimation: Bounded Response

We tested the ability of the beta mixed-effects model to accurately estimate relationships among variables. This relationship is linear on the scale of the link function. Due to the effect of the link function in beta regression, the parameters in that model are not directly comparable to those in the various linear regressions tested thus far, so the comparison here is between beta regression models that do and do not take non-independence among the observations into account via the inclusion of the random effects term. We fit two models to each simulated dataset: a beta mixed-effects model (equation 8) and a standard beta regression model with no random effects term (equation 6). Fig. 7 shows the comparison of their parameter estimation. To better visualize the impact of data non-independence, we also fit standard beta regression models to datasets simulated with no random effects and with random effects that had no dependency structure (i.e. 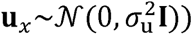.

**Figure 7.**
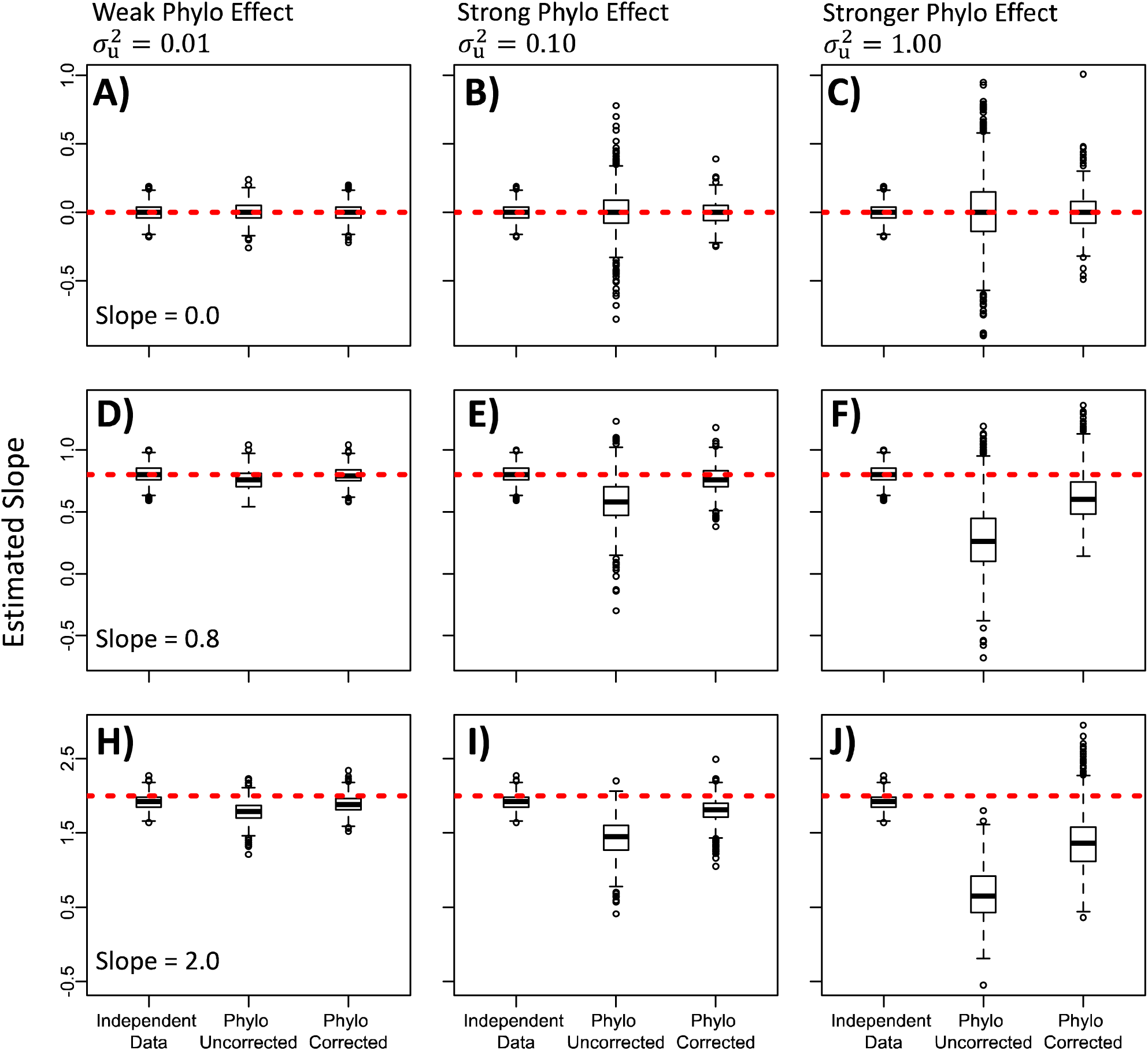
Parameter estimation in beta regression. Values are replicate means of posterior distributions for Bayesian-estimated regression coefficients. The leftmost boxplot in each panel shows the posterior means estimated when fitting a standard beta regression model to datasets simulated with no dependency structure (this boxplot is identical for the three panels in each row); middle and right boxplots show results from fitting that same beta regression model and the beta mixed-effects model, respectively, to datasets simulated with lineage-pair non-independence. Dashed red lines show the simulated slopes of the relationships on the scale of the link function; slopes apply to their entire row. Each model was fit to 1,000 simulated datasets with bounded responses. All models were given the same prior distributions of *b*_1_ ∼N(0,10) and assumed a logit link function. The 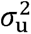 parameter scales the random effects: higher values indicate greater influence of non-independent random effects on the response variable.

Our simulations show that it is critical to take non-independence into account when using beta regression to analyze bounded lineage-pair data. Comparison of the left and middle boxplots in each panel of Fig. 7 shows that lineage-pair non-independence has a substantial effect on accuracy and precision of parameter estimation. Comparison of the middle and right boxplots show that accounting for non-independence via the mixed-effects term recovers much of that accuracy and precision: the beta mixed-effects model outperforms a beta regression model across all parameter combinations. Our results also show than when the simulated effect of non-independence is extreme and regression coefficients are nonzero, even the substantial improvements provided by the beta mixed-model do not fully recover lost accuracy; in such cases, we detect a slight downward bias in estimates of the slope parameter (on the scale of the link function) from the mixed-effects model (Fig. 7C,F,J), and this is more pronounced when the simulated linear relationship is severe. Such a strong linear relationship induces a slight downward bias even when fitting a standard beta regression model to independent data (see left boxplots in Fig.7H-J). However, even in the most severe scenario (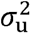 = 1), the correct positive relationships were returned (i.e. Bayesian posterior 95% credible intervals did not include zero) in 99.1% and 100% of estimates from beta mixed-effects models with nonzero slopes (*b*_1_ = 0.8 and *b*_1_ = 2.0, respectively).

### Empirical Applications

We illustrate the empirical application of our new approaches by comparing the fit of several models to two published datasets. The first is the previously mentioned Price and Bouvier (2002) bird hybridization dataset. The second is a dataset of sexual isolation (based on metrics in Sobel and Chen 2014) and ‘ecological distance’ (based on Funk et al. 2006) in allopatric pairs of *Drosophila*, which we obtained from the drosophila-speciation-patterns.com website hosted and curated by Roman Yukilevich (data file included as supplementary material). Our intention is to provide a practical example of model fitting rather than to comprehensively test hypotheses, so we applied coarse filters when data wrangling and made no effort to update the taxonomy in the trees or corresponding the data tables. We evaluated model support based on leave-one-out cross validation (LOO) and the Watanabe-Akaike or “widely applicable” information criterion (WAIC) (Vehtari et al. 2017, Watanabe 2010, Hooten and Hobbs 2015).

Results from bird hybridization data show that accounting for non-independence via the **C***_P_* matrix has a powerful effect on both model support and parameter estimation. The linear-pair LMM is strongly supported over OLS and Castillo’s (2017) two-term LMM (the other two linear regression models fit to original dataset) with vastly lower scores for both LOO and WAIC (Table 2). Accounting for non-independence with **C***_P_* also nearly doubled the strength of the relationship between divergence time and hybrid sterility: slopes in OLS and the linage-pair mixed model were 0.08 and 0.15, respectively. We found similar patterns among beta regression models fit to transformed data. The beta mixed model is strongly supported over standard beta regression and yields more than a two-times stronger relationship between divergence time and hybrid sterility (Table 2). In all models, the credible intervals for the slope parameter did not include zero.

Accounting for non-independence via the **C***_P_* matrix had a lesser impact when analyzing the *Drosophila* dataset, but results from *Drosophila* show that model support was profoundly improved by using beta regression compared to all forms of linear regression, highlighting the utility of this framework for analyzing bounded biological data (Geissinger et al. 2022). Contrary to the bird results, our mixed-effect models using **C***_P_* in both linear and beta regression applications had negligible impacts on model support or slope parameter estimation. A potential explanation is that experimental fly crosses were primarily conducted within (rather than among) various groups of closely related species. It is possible that close relatives indeed covary in the extent of their sexual isolation with other taxa, but much of that sexual isolation went unmeasured, thus the underlying covariance is not reflected in the dataset. The credible intervals for the slope parameter did not include zero for any models fit to these *Drosophila* data, suggesting confidence in the relationship between the two sampled variables.

### Conclusion

Relationships among pairwise-defined traits are of great potential interest to biologists working in numerous subfields. The comparative analysis of these lineage-pair data has already played an important role in certain areas of evolutionary biology, in particular the field of speciation research, but the broader adoption of this approach has been hindered by a lack of clarity in terms of analytical approach and a lack of ready-made tools for empiricists to employ. Our first goal in this paper was to show one way to deal with the inherent dependency of lineage-pair data. We have demonstrated that the hypothesized covariance structure of lineage-pair traits can be obtained from a phylogeny by making some common and simple assumptions about how traits evolve within species and about how those traits affect pairwise-defined characters of interest. These assumptions imply models from which emerge the expected covariance structure of lineage-pair traits, which can be used to account for non-independence in regression-like analyses. Our second aim was to provide a dedicated computational tool, the R package *phylopairs*, that will permit empiricists to begin to address a diversity of interesting questions based on this framework and that will provide them some flexibility in their choice of approach. Our final goal for this paper is to help open the door to the creative development of comparative analyses based on lineage-pair data. We hope that by helping to clarify the expected covariance structure of species-pair traits in the continuous case, and under simple Brownian motion assumptions, we will spur others to pursue a variety of improvements and elaborations on this basic framework.

## Materials and Methods

### Parameter estimation: unbounded response

We simulated 1,000 pure-birth phylogenetic trees of 20 species and simulated a continuous trait evolving under Brownian motion through each tree. For each of the 1,000 replicates, we made a table with 190 rows, each row containing one unique pairwise combination of the 20 species in the tree along with the square difference between those species in their simulated trait. We next generated a single predictor variable for each replicate by randomly drawing 190 observations from a standard normal distribution. We defined an intercept and a slope and calculated a deterministic response variable as a linear function of the predictor. To this deterministic response we added two sources of variation. The first was the squared differences between species in their simulated trait, scaled by a parameter 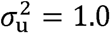. The second was a set of uncorrelated residuals drawn from a multivariate normal distribution with variance 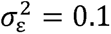. All simulations were conducted in R version 4.3.2 and used functions from the *ape*, *phytools*, and *MASS* packages (Paradis and Schliep 2019, Revell 2024, Venables and Ripley 2002), in addition to functions from *phylopairs*.

The simulation procedure described above follows a model in which the pairwise-defined response variable is linearly related to the square difference between species in a trait evolving under BM. Both the unweighted and weighted node averaging procedures make no explicit assumption about an underlying model of evolution, so their performance should not be affected by this choice. Castillo’s modified PLMM likely implies an underlying model of evolution, but it is not obvious what that model is. We note that it would have been possible to simply add the random effects in our simulation by drawing from a multivariate normal distribution with mean 0 and variance 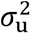, but as this is the exact structure of our mixed effects LMM, such an approach might overly advantage that method.

To each replicate dataset we fit our lineage-pair LMM, the two-term LMM, a standard OLS regression model, and both node averaging procedures. For the latter, we first calculated the average of the predictor and response variables at each node of the tree as described by Coyne and Orr (1989, unweighted) and Fitzpatrick (2002, weighted) and then fit an OLS model to the resulting values. The fitting of the two-term LMM further requires the calculation of two design matrices that map the correct “species 1” and “species 2” random effect to each pair (Castillo 2017). We conducted all simulation and model-fitting procedures twice, once with zero slope (*b*_1_ = 0) and once with a positive slope of *b*_1_ = 0.8; in both cases the intercept was arbitrarily set to *b*_0_ = 1. We fit all models in a Bayesian context via the *phylopairs* package.

### Sensitivity to Missing Data

We simulated 500 replicate datasets at each of five levels of missing data. It is important to ensure that the final datasets are the same size at each level so that changes in model performance do not simply result from ever-decreasing statistical power. Our approach was to first stipulate that each final dataset will have 30 lineage pairs (but note that node averaging methods reduce this number further). Next, we determined how many taxa to include in the phylogenetic trees such that 30 pairs would represent different percentages of missing data. Combinatorics work with whole numbers, so it is not possible to specify *a priori* that 30 pairs will represent, say, 50% of the total pairs that can be made from the species in a tree, because no whole number of taxa will combine to make exactly 60 pairs. After trial and error, we ended up with five tree sizes for which 30 pairs comprise a good range of proportions of missing data. The number of species and corresponding proportions of missing data at each of the five levels are shown in Table 1. The smallest proportion of missing pairs was 16.67%; the largest was 94.06%.

**Table 1.** Relationship between divergence time and reduction of hybrid fertility in birds.

|  | Estimated Slope | LOO | WAIC |
| --- | --- | --- | --- |
| OLS | 0.08 | 342.2 | 342.2 |
| LP-LMM | 0.15 | 136.8 | 64.8 |
| TT-LMM | 0.08 | 321.8 | 303.2 |
| Beta Regression | 0.06* | -13.8<sup>†</sup> | -13.9<sup>†</sup> |
| Beta Mixed Model | 0.15* | -140.8<sup>†</sup> | -196.4<sup>†</sup> |

Lower values of both LOO and WAIC denote better model fit. Models with greatest support for their respective data types are highlighted in grey.
\* This is the slope of the predicted value of the response variable on the scale of the link function.
<sup>†</sup> Support for beta regression models is not directly comparable with that of the linear regressions in this table as the two groups of models were fit to response variables with different bounds.

**Table 2.** Estimating relationship between ecological distance and sexual isolation in allopatric *Drosophila*.

|  | Estimated Slope | LOO | WAIC |
| --- | --- | --- | --- |
| OLS | 0.27 | 32.9 | 32.9 |
| LP-LMM | 0.28 | 32.9 | 32.6 |
| TT-LMM | 0.17 | 28.2 | 23.1 |
| Beta Regression | 0.92* | -49.4 | -49.4 |
| Beta Mixed Model | 0.92* | -49.3 | -49.3 |

Lower values of both LOO and WAIC denote better model fit. \* This is the slope of the predicted value of the response variable on the scale of the link function.

We first simulated a series of complete datasets. We generated 500 pure-birth phylogenetic trees of each tree size shown in Table 3. We then repeated the exact simulation procedure from the previous test to end up with observations for the predictor and response variables for every possible pair that could be constructed from each tree in each replicate. The linear relationship between response and predictor was fixed at *b*_0_ =1 and *b*_1_ = 0.8.

**Table 3.** Numbers of species and possible pairs at 5 levels of data incompleteness.

|  | Number of<br>Taxa in Tree | Number of<br>Possible Pairs | Proportion of<br>Pairs Missing* |
| --- | --- | --- | --- |
| Level 1 | 9 | 36 | 16.67 |
| Level 2 | 10 | 45 | 33.33 |
| Level 3 | 12 | 66 | 54.54 |
| Level 4 | 16 | 120 | 75.00 |
| Level 5 | 35 | 595 | 94.06 |

\*Calculated based on a final dataset of 30 lineage pairs

We then generated incomplete datasets by randomly selecting 30 pairs from each data table at each treatment level and discarding all other pairs. The removal of pairs affects the size of **C***_P_*; all final **C***_P_* matrices had dimensions 30 × 30. The size of the **C** matrices in Castillo’s two-term LMM is only affected if the total number of species included in the analysis changes (often a large number of pairs can be removed without affecting the total number of species in a dataset). Whether or not the values in **C***_P_* and c are altered by the removal of pairs depends on whether the total number of species remaining in the dataset changes and whether trees are pruned prior to calculating covariance matrices. When the total number of species is the same, the expected covariances among pairs in **C***_P_* and among species in c is the same. If there are fewer total species in the dataset after removing some pairs, then the covariances among species and pairs might change *if* the phylogenetic tree is pruned to include just the remaining taxa prior to calculating covariances. For consistency, we did not recalculate covariances in either **C***_P_* or c but simply removed the rows and columns corresponding to missing pairs or species, respectively. We did recalculate the design matrices for Castillo’s PLMM to ensure that random effects properly mapped to each pair.

Two consequences arise from this simulation procedure. First, the size of incomplete datasets for the node averaging methods were not constant within and across treatments (unlike for the lineage-pair LMM, Castillo’s PLMM, and OLS). “Incomplete” node averaged datasets at Level 1 ranged in size from 5 to 8 values with a median of 8, and those at level 5 ranged from 5 to 15 values with a median of 9. This variability is unavoidable, and its source is described in detail in the Supplementary Text. Second, while the identity of missing pairs is entirely random in our simulation, this will not always be the case in empirical studies. If an empirical response variable is something like hybrid growth rate, for example, then there may be a bias such that pairs comprised of more deeply diverged taxa are more likely to be excluded from the analysis (i.e., if such crosses are more likely to fail).

We fit each of the five models as previously described to each complete and incomplete dataset. We then calculated the percentage difference in the coefficient estimated from each incomplete dataset relative to the complete datasets from which it was derived.

### Bias in Weighted Node Averaging due to Missing Data

To further examine the apparent underestimate in weighted node-averaged values observed in the ‘sensitivity to missing data’ tests, we first generated a single phylogenetic tree with 50 tips (49 nodes). We then generated 1,000 replicate data tables of 1,225 rows corresponding to the number of pairs that can be derived from 50 taxa. We simulated an arbitrary positive-valued pairwise-defined trait by taking 1,225 random draws from a half-normal distribution for each replicate table. We then made 1000 incomplete data tables by randomly removing a pre-defined proportion of rows from each dataset; this procedure was repeated for six proportions of missing data (∼ 5%, ∼15%, ∼25%, ∼50%, ∼75%, and ∼90%). We fit OLS models to weighted and unweighted node averages and calculated error as the percent difference between node averages as calculated from incomplete versus complete datasets, as shown in Fig. 6.

### Parameter Estimation: Bounded Response

We simulated linear relationships among variables on the link scale and then converted the response variable to the beta distribution via logistic transformation. Our simulations thus assume a logit link function. To simulate 1000 replicate linear predictors ***η***, where ***η* = X*β* + u**, we first generated replicate vectors of a single predictor variable by taking 1000 random draws of 190 values from a standard normal. Each vector becomes the second column in an **X** matrix, where the first column is 1s and corresponds to the intercept (which we arbitrarily set to *b*_0_ = 0). To generate **u** vectors, we simulated 1000 pure-birth trees of 20 taxa and calculated the linear-pair covariance matrices **C***_P_* from each tree. We then randomly drew 1000 sets of 190 values from multivariate normal distributions (where 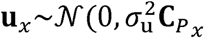) for the *x^th^* replicate). After adding the replicate **X*β***s and **u**s to get replicate ***η*** vectors, we next calculated the predicted values of the responses in each replicate by taking the logistic function of the linear predictor as 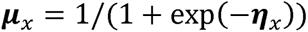. We fixed the precision parameter at **φ** = 5. From our replicate vectors of ***µ*** and the fixed value of **φ**, we calculated replicate sets of the two shape parameters *p* and *q* that the R software uses to define the beta distribution:

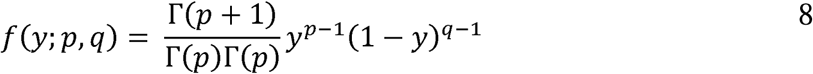

where *p* = *µ*/( *µ* + 1) and *q* = **φ** − *p*. We finally generated the response variables via random draws from beta distributions defined by those shape parameters. We conducted the above procedure for nine parameter combinations corresponding to three scales of random effects 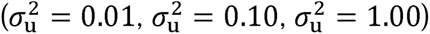 and three slopes 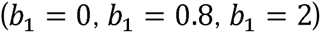.

### Empirical Analyses

Both analyses required a dataset and a phylogenetic tree. We obtained a dated maximum clade credibility tree for all birds from Pulido-Santacruz and Weir (2016), which combines two earlier trees (Jetz et al. 2012; Derrberry et al. 2011). For flies, we used a new drosophilid tree from Kim et al. (2023) that has 365 species including 279 from the genus *Drosophila*. We converted the undated Kim et al. (2023) tree to a relative time ultrametric tree via the ‘chronos’ function from the ‘ape’ package (Paradis and Schliep 2019). We found that a lambda smoothing parameter of zero was best supported by likelihood. We trimmed each dataset down to the pairs whose species were both represented in the respective tree. This eliminated all crosses between subspecies and geographic races, which were abundant in the two datasets. The final bird and *Drosophila* datasets contained 111 and 61 pairs, respectively. The response variable for *Drosophila* was bounded between 0 and 1, while in birds it was bounded between 1 and 5. For the bird dataset we fit the linear models to the original data and fit beta regression models to a transformed response (where the response was simply normalized by dividing by 5). Support for the linear and beta regression models is thus not directly comparable in the case of bird data (Table 2).

## Acknowledgements

Thanks to Joe Felsenstein, Matt Pennell, James Boyko, Andrius Dagilis, and Ken Thompson for their time and thoughtful consideration. We are grateful to Michael Turrelli, Yaniv Bradvain, and Roman Yukilevich for facilitating access to their data and/or code. Thanks to members of the Matute lab for comments on the manuscript. This work was supported by the National Institute of General Medical Sciences under Award R35GM148244.

## Appendix 1

We model an underlying biological character, *X*, as a continuous trait that evolves via Brownian motion (BM) independently in each branch of a phylogenetic tree. We set the ancestral mean value of *X* at the root of the tree to zero, which means that if species *i*, *j*, *k*, and *l* are species in the tree, then *X_i_*, *X_j_*, *X_k_*, and X*_l_* are jointly distributed zero-mean Gaussian variables with phylogenetic covariance matrix **C**. This matrix is scaled by the BM dispersion parameter 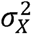.

We assume that the pairwise-defined response variable of interest is linearly related to a random variable *Y*, where *Y* can be related to *X* by one of three relationships.

### Relationship 1: *Y_ij_* = (*X_i_* − *X_j_*)^2^

If *Y* is the squared difference between species in *X*, where *Y_ij_* = (*X_i_* − *X_j_*)^2^, then we describe the covariance among pairs in *Y* in terms of the covariance among species in *X* as follows.

If we have two species pairs, one comprised of species *i* and *j* and the other of species *k* and *l*, then the between-pair covariance of the pairwise-defined traits *Y_ij_* and *Y_kl_* is defined by

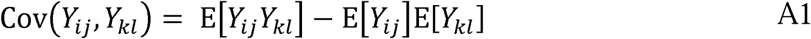

We expand the second term on the right side of A1 to get E[*Y_ij_*] = E[(*X_i_* − *X_j_*)^2^] = E[*X_i_*^2^] − 2E[*X_i_X_j_*] + E[*X_j_*^2^]. From the definition of the variance of a random variable, E[*X_i_*^2^] = Var(*X_i_*) − E[*X_i_*]^2^, and since E[*X_i_*] = 0, this becomes 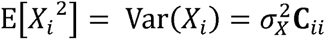. Likewise, 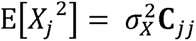. From the definition of covariance, E[*X_i_ X_j_*] = Cov(*X_i_ X_j_*) + E[*X_i_*]E[*X_j_*], and since E[*X_i_*] = E[*X_j_*] = 0, we have 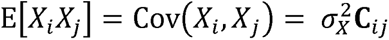. Thus, 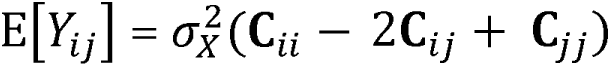. Likewise, 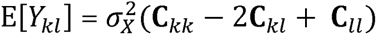. The product being subtracted as the second term of equation A1 is then

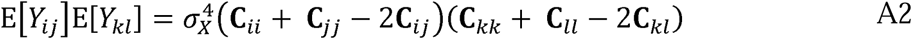

Now considering the first term on the right side of A1, we expand E[*Y_ij_ Y_kl_*] = E[(*X_i_* − *X_j_*)^2^(*X_k_* − *X_l_*)^2^] to get

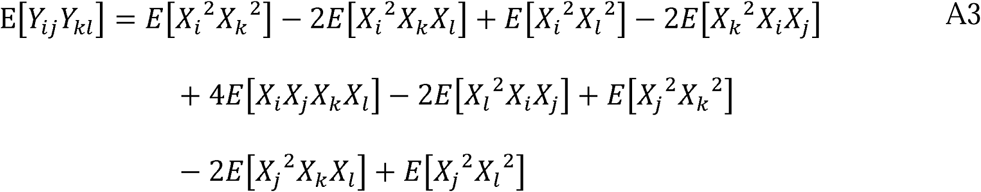

Each term is a higher-order moment of a zero-mean multivariate normal distribution. We can use Isserlis’ theorem to express each higher-order moment in terms of the covariances of the variables involved. By Isserlis’ theorem in the even case, we first express E[*X_i_ X_j_ X_k_ X_l_*] as

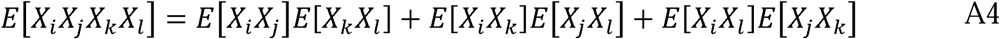

We have shown above that when E[*X_i_*] = E[X_j_] = 0, then 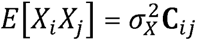, so A4 simplifies to

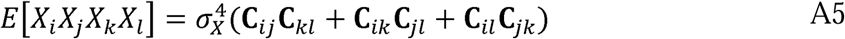

Each additional term in A3 can be written in the form of E[*X_i_ X_j_ X_k_ X_l_*] (e.g. E[*X_i_*^2^*X_k_*^2^] can be written as E[*X_i_ X_j_ X_k_ X_l_*], and E[*X_i_*^2^ *X_k_ X_l_*] can be written as E[*X_i_ X_j_ X_k_ X_l_*]). Applying Isserlis’ theorem in the exact same way as above for every additional term in A3 and adding the terms gives us the first term on the right side of equation A1:

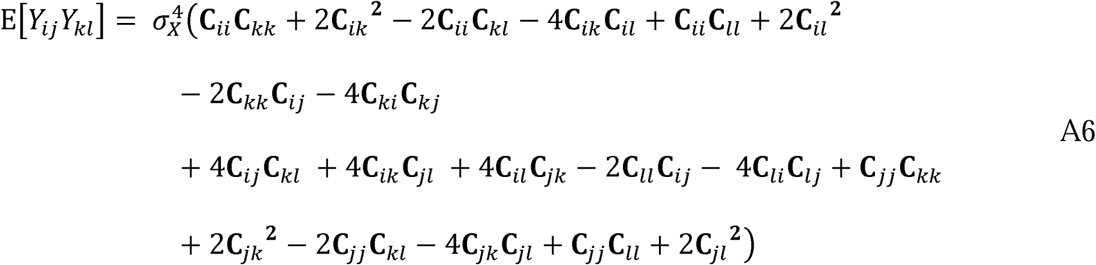

Subtracting A2 from A6 in accordance with A1, we finally get

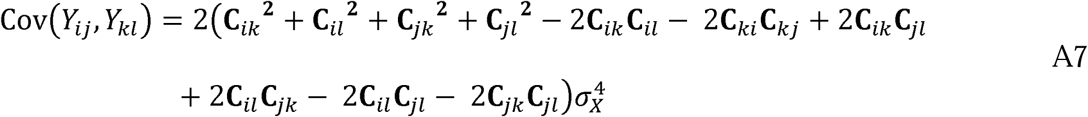

We assume that our pairwise-defined trait of interest is linearly related to *Y*, which means that the expected covariance in this trait among pairs is equivalent to A7 but with a unique scale parameter. The unscaled version of the covariance is then used in downstream applications, and the scale is estimated from the data. Note that the covariance matrix so-produced is often not well-conditioned, but statistical analyses are nonetheless substantially improved by its use compared to alternative approaches.

### Relationship 2: *Y_ij_* = (*X_i_ + X_j_*)^2^

Following the steps above, the unscaled covariance when *Y_ij_* is the squared sum of *X_i_* and *X_j_* is

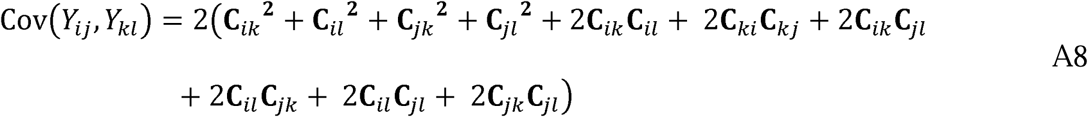

### Relationship 3: *Y_ij_ = X_i_ X_j_*

Following the steps above, the unscaled covariance when *Y_ij_* is the product of *X_i_* and *X_j_* is

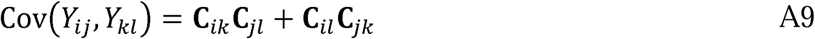

## Supplementary Text

### Variability in “Incomplete” Dataset Size for Node Averaging Procedures

The two node averaging procedures were unique in that the size of the incomplete datasets they generate were not constant across treatments in our ‘missing’ data simulations. This is an unavoidable consequence of how node averaged values are created.

Recall that the maximum size for these datasets is equal to the number of nodes in a trees, which is *N*- 1 in a fully bifurcating tree with *N* tips. At Level 1 of our simulation, all initial trees contained 9 taxa and 8 internal nodes. There are 36 possible pairs that can be created from these taxa. After randomly removing 6 pairs to create a final dataset of size 30, it is possible (but unlikely) that, by chance alone, some nodes are no longer spanned by two species comprising a remaining pair. In such a case, those nodes will drop out, and the final node-averaged dataset will be smaller than its maximums size of 8. By contrast, at Level 5 of our simulation, the number of nodes in the tree is 34. Since the number of nodes in the tree is greater than the number of pairs in the final dataset (30), it is now inevitable that many nodes will no longer be spanned by two species that comprise a remaining pair. The number of those 34 nodes that are still spanned by species in a final dataset pair will vary by chance across replicates. Note that we repeated all node average analyses using an approach in which we first pruned the original species tree to contain just the taxa included in the final dataset. This pruning had no effect on final dataset sizes. The reason is that at higher treatment levels, the number of species included among the final 30 pairs tended to be greater, since more species were included in the full datasets and we drew from those datasets randomly. This means that the size of the trees generated from the final datasets were still larger at higher treatment levels, despite our pruning. Those larger trees have more internal nodes, and which of those nodes are spanned by taxa in the final dataset is still a matter of chance.

**Figure S1.**
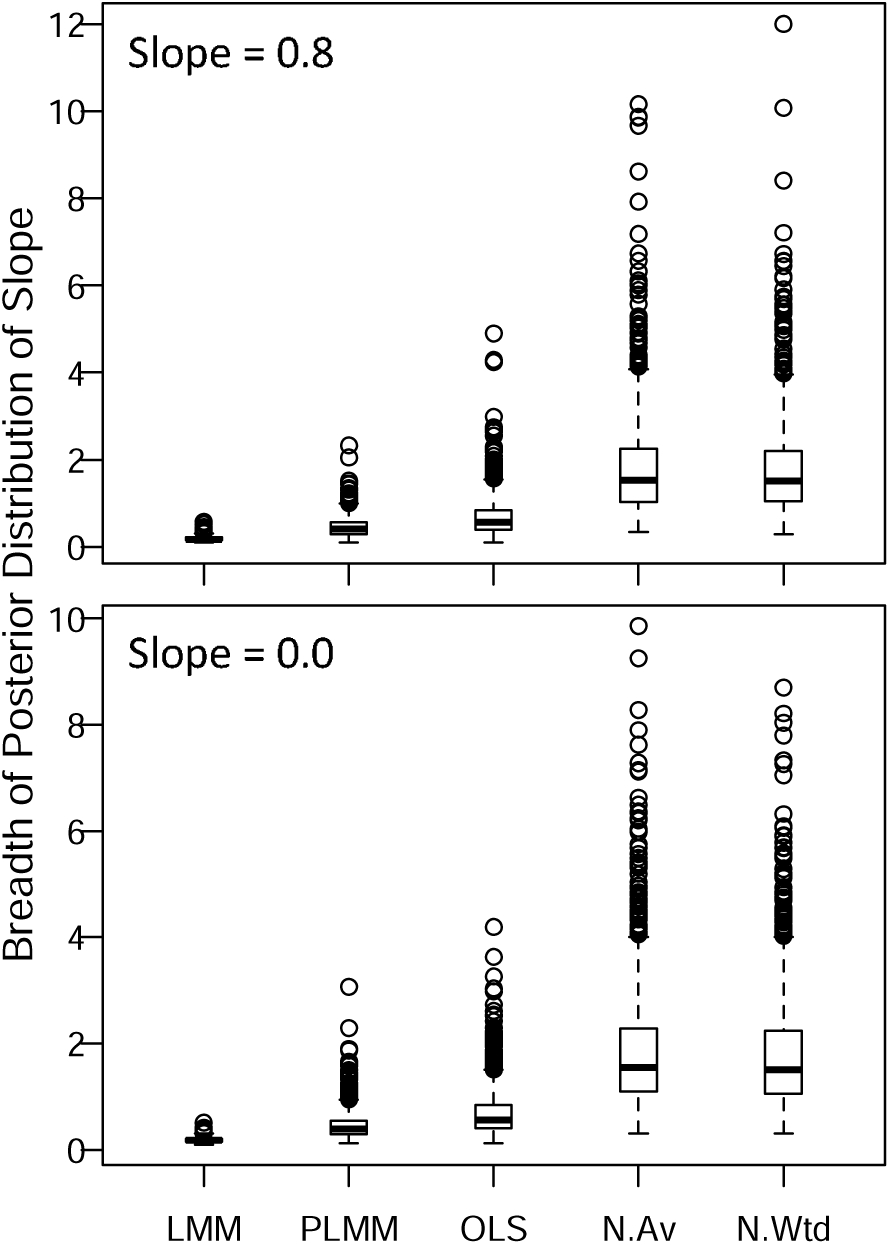
Breadth of posterior distributions of slope estimates. Values are 95% credible intervals of the posterior distributions of the regression coefficient. Each model was fit to 1000 simulated datasets with unbounded responses. “N.Wtd” and “N.Av” indicate weighted and unweighted node averaging, respectively. The lineage-pair LMM, the two-term LMM, and OLS models were fit to datasets with 190 observations. Node averaging condenses these datasets to the number of nodes in the tree (19). Note that Y-axes are not equivalent.

