## Supplementary data and code for "The comparative analysis of lineage-pair traits": anderson.etal.analyses.R

```
#======================================================
## CODE FOR THE 5 ANALYSES IN ANDERSON ET AL. 2024 (SUBMITTED)
#======================================================
# The code in this file details how each of the three model performance analyses and two empirical analyses were run.
# The three performance analyses were run on the "Longleaf" cluster at UNC Chapel Hill.
# The code below for generating simulated datasets is exactly as was used for the paper.
# The code below for fitting models is the unscaled version of the code used for the paper.
# That is, the code below shows how to fit models to the dataset, but fitting the models to all simulated data will typically require a cluster, and the correct way to scale this code for cluster use will depend on the cluster
# We leave it to the user to adapt the model performance model-fitting code shown here as appropriate for their computational approach and resources.  
# For the underlying code of the 'phylopairs' package funtions, see package materials on the package's CRAN page. 
# Code here and 'phylopairs' package written by Sean A. S. Anderson
# The five analyses are: 
# Analysis 1: Parameter estimation, unbounded response variable
# Analysis 2: Effect of missing data on parameter estimation
# Analysis 3: Parameter estimation, bounded response variable
# Analysis 4: Re-analysis of empirical Drosophila data from
# Analysis 5: Re-analysis of empirical avian hybridization data from Price and Bouvier (2002)
# The code here is not written to be run as a script and code from different analyses is no meant to be run in a single session 
# (some variable names are re-used in different Analysis blocks; packages are re-loaded and functions re-defined in each Analysis block as necessary)
#======================================================


#======================================================
## Analysis 1: Parameter Estimation, Unbounded Response
#======================================================

##===================
## A) Simulating Replicate Datasets of Unbounded Pairwise-Defined Traits
##===================
library(phytools)
library(MASS)
library(phylopairs)
Nsim=1000
Nsp=20
# Generate replicate trees of 20 taxa
trees=pbtree(n=Nsp, nsim=Nsim)
# Make sure they're all ultrametric
all(sapply(trees, is.ultrametric))
# Generate a set of traits evolving under Brownian motion (BM) for each replicate tree
X=lapply(trees, fastBM)
# For each replicate trait dataset, find the pairwise combination of pairs
xps = lapply(trees, function(x) data.frame(t(combn(x$tip.label,2))))
for(i in 1:length(xps)) {
  colnames(xps[[i]]) = c("sp1","sp2")
  xps[[i]] = cbind(xps[[i]], t(combn(X[[i]],2)))
}
# Ensure that each replicate has the same number of species pairs
length(unique(lapply(xps, nrow))) #1
# Calculate Y, the square difference in the BM trait values for each pair
ypairs = t(sapply(xps, function(x) (x[,3]-x[,4])^2)) # each row is a replicate vector of Y=(Xi-Xj)^2
# Calculate the covariance matrix among pairs in Y for each replicate (takes a few minutes)
cov.y=list()
for(i in 1:Nsim) cov.y[[i]] = taxapair.vcv(sp.pairs=xps[[i]], tree=trees[[i]], spnames=TRUE)
# Calculate the Z matrices and covariance matrices for each replicate dataset (required for Castillo method only, which we refer to as the two-term plmm)
# There are two Z matrices that map, respectively, the species1 and species2 random effects to each observation
# There are two phylogenetic covariance matrices that define, respectively, non-independence within species1 and species2 random effects
# Note that not all species necessarily will be 'species 1' and 'species 2' in a given dataset
# The function twoterm.plmm.mats takes this into account when generating the four matrices
re.mats=list()
for(i in 1:Nsim) re.mats[[i]] = twoterm.plmm.mats(sp.pairs=xps[[i]], tree=trees[[i]])
# Verify that the covariance matrices are valid using the covmat.check function
all(sapply(re.mats, function(x) sapply(x[3:4], function(y) all(covmat.check(y)))))
all(sapply(cov.y, function(x) all(covmat.check(x))))
# Generate a predictor variable with draws from standard normal
pred=t(replicate(Nsim, expr=rnorm(nrow(xps[[1]]))))
# Put each set of predictor variables into a design matrix and join these matrices in a list
xmats = apply(pred, 1, function(x) cbind(rep(1, length(x)), x), simplify=F)

### FOR ANALYSIS 1 AND 3, THE ABOVE PROCEDURE IS ONLY CONDUCTED ONCE
### VARIATION IN THE SIMULATION PROCEDURES (I.E. DIFF REGRESSION PARAMETERS, BOUNDED VERSUS UNBOUNDED RESPONSE) CAN BE ADDED DOWNSTREAM STARTING NOW

# Define an intercept and slope for the relationship between response and predictor
b0=1
b1=0.8
# Calculate predicted val of response variable based on predictor of each rep
resp.x = b0 + b1*pred
# Calculate and add structured random effects and residuals to response
# First, generate, for each rep, random effects that are scaled versions of Y (a*Y, where Y=(xi-xj)^2)
# For simplicity, we set a=1
sig2.rand = 1
u = ypairs*sig2.rand
# Second, generate, for each rep, actual random error
sig2.eps=0.1
eps = mvrnorm(n=Nsim, mu=rep(0, nrow(cov.y[[1]])), Sigma=diag(nrow(cov.y[[1]]))*sig2.eps)
# Compile these effects together to get an 'observed' response variable
resp = resp.x + u + eps
# Visualize the relationship between response and predictor
plot(pred[5,], resp[5,])

# Now, we can use the node averaging functions of phylopairs to condense these datasets to just the node-averaged values
# We need to condense both the predictor variables and the response variables
# We need to do this both for the simple node averaging procedure and the weighted node averaging procedure
# For these functions, the separate components of the datasets need to be joined into a single tabular dataset
node.av=node.av.wtd=list()
for(i in 1:Nsim) {
	set = cbind(xps[[i]][,1:2], pred[i,], resp[i,])
	pred.nodes=node.averager(tree=trees[[i]], taxacolumns=c("sp1","sp2"), dataset=set, varb="pred[i, ]")
	resp.nodes=node.averager(tree=trees[[i]], taxacolumns=c("sp1","sp2"), dataset=set, varb="resp[i, ]")
	pred.nodes.wtd=node.averager(tree=trees[[i]], taxacolumns=c("sp1","sp2"),dataset=set, varb="pred[i, ]", weighted=TRUE)
	resp.nodes.wtd=node.averager(tree=trees[[i]], taxacolumns=c("sp1","sp2"),dataset=set, varb="resp[i, ]", weighted=TRUE)
	node.av.mat=cbind(rep(1, length(pred.nodes)),pred.nodes)
	node.av.wtd.mat=cbind(rep(1, length(pred.nodes.wtd)),pred.nodes.wtd)
	node.av[[i]]=list(node.av.mat, resp.nodes)
	node.av.wtd[[i]]=list(node.av.wtd.mat,resp.nodes.wtd)
}

# Save the predictor variable design matrices, response variables, phylogenetic covariance matrices, and Z matrices as an R data object
save(trees, xmats, resp, cov.y, re.mats, xps, ypairs, node.av, node.av.wtd, file="data.to.fit.unbounded.rdata")
# We repeated the above procedure but with b1=0 and saved the resulting objects as "data.to.fit.unbounded.zeroslope.rdata"
##===================

##===================
## B) Fitting Models to Simulated Datasets 
##===================
# Fitting models to all replicate simulated datasets within a reasonable time will typically require the use of a compute cluster
# Computing clusters vary in terms of how replicate jobs are run, so the code below is for fitting a model to a single replicate dataset
# This code can be modified and incorporated into scripts for fitting models to multiple replicate datsets in parallel
# For model-performance testing, we set iter=6000 for each model fitting function (where iter is the number of sampling iterations in the underlying Bayesian models)
library(phylopairs)
# Load dataset
load("data.to.fit.unbounded.rdata")
# Define which replicate to run 
num=1
# Define the number of iterations to run
its=6000
# Fit OLS 
outs.ols = linreg.stan(des=xmats[[num]], y=resp[num,], model="ols", iter=its)
# Fit the new lineage-pair LMM
outs.lplmm = linreg.stan(des=xmats[[num]], y=resp[num,], model="pgls.mm", covmat = cov.y[[num]], iter=its)
# Fit Castillo's (2017) LMM
outs.lmm = twoterm.plmm.stan(des=xmats[[num]], y=resp[num,], sp1=xps[[num]]$sp1, sp2=xps[[num]]$sp2, tree=trees[[num]], iter=its)
# Fit OLS to unweighted node averages
outs.node = linreg.stan(des=node.av[[num]][[1]][,2], y=node.av[[num]][[2]], model="ols", iter=its)
# fit OLS to weighted node averages
outs.node.wtd = linreg.stan(des=node.av.wtd[[num]][[1]][,2], y=node.av.wtd[[num]][[2]], model="ols", iter=its)
# Observe parameter estimates (slope estimate is coef2)
outs.ols[[1]]
outs.lplmm[[1]]
outs.lmm[[1]]
outs.nodav[[1]]
outs.nodav.wtd[[1]]
##===================

##===================
## C) Sumarizing Results
##===================
# Here we show how we summarized the results and made the graphs that form the basis for fig4. 
# In the interest of compute time, and for illustrative purposes, we show this for three replicate runs of each model and with only 1000 Bayesian iterations; the actual graphs in Fig4 are based on 1000 replicate runs each with 6000 Bayesian iterations.
ols.slopes = lplmm.slopes = lmm.slopes = node.slopes = node.wtd.slopes = rep(NA, 3)
its=1000
for(i in 1:3) {
	ols.slopes[[i]] = linreg.stan(des=xmats[[i]], y=resp[i,], model="ols", iter=its)[[1]][2,"mean"]
	lplmm.slopes[[i]] = linreg.stan(des=xmats[[i]], y=resp[i,], model="pgls.mm", covmat = cov.y[[i]], iter=its)[[1]][2,"mean"]
	lmm.slopes[[i]] = twoterm.plmm.stan(des=xmats[[i]], y=resp[i,], sp1=xps[[i]]$sp1, sp2=xps[[i]]$sp2, tree=trees[[i]], iter=its)[[1]][2,"mean"]
	node.slopes[[i]] = linreg.stan(des=node.av[[i]][[1]], y=node.av[[i]][[2]], model="ols", iter=its)[[1]][2,"mean"]
	node.wtd.slopes[[i]] = linreg.stan(des=node.av.wtd[[i]][[1]], y=node.av.wtd[[i]][[2]], model="ols", iter=its)[[1]][2,"mean"]
}
# Compare each method to OLS 
reps = 3
lplmm.comp = lmm.comp = node.comp = node.wtd.comp = rep(NA, 3)
for(i in 1:reps) {
  if(!is.na(lplmm.slopes[i])) {
    if(abs(0.8 - lplmm.slopes[i]) < abs(0.8 - ols.slopes[i])) lplmm.comp[i]=1
  }
  if(abs(0.8 - lmm.slopes[i]) < abs(0.8 - ols.slopes[i])) lmm.comp[i]=1
  if(abs(0.8 - node.slopes[i]) < abs(0.8 - ols.slopes[i])) node.comp[i]=1
  if(abs(0.8 - node.wtd.slopes[i]) < abs(0.8 - ols.slopes[i])) node.wtd.comp[i]=1
}
lplmm.win=length(which(lplmm.comp==1))/reps
lmm.win=length(which(lmm.comp==1))/reps
node.win=length(which(node.comp==1))/reps
node.wtd.win=length(which(node.wtd.comp==1))/reps
# Calculate Relative Wins of Each Procedure (win = estimate closer to simulated value)
correction = c(lplmm.win, lmm.win, node.win, node.wtd.win)
ols = 1-correction
rel.wins = rbind(correction,ols)

## Graph the Results
# Parameter Estimates
boxplot(list(lplmm.slopes, lmm.slopes, ols.slopes, node.slopes, node.wtd.slopes), col=rgb(0,0,0,0), boxwex=.25, yaxt='n', ylim=c(-2, 3.5), names=c("LMM", "PLMM", "OLS", "N.Av", "N.Wtd"))
axis(2, labels = NA, tcl=-0.3)
axis(2, lwd = 0, line = -0.6, las=1)
abline(h=0.8, col="red", lwd=3, lty=3)
mtext("Estimated Slope", side = 2, line = -1.5, outer = TRUE, las=0, cex=1.2)
# Relative Wins
colnames(rel.wins)=c("LMM", "PLMM", "N.Av", "N.Wtd")
barplot(rel.wins, col=c("grey", "white"), yaxt='n', width=rep(0.4,4))
legend(x=par("usr")[1], y=par("usr")[4] * 1.15, legend=c("Correction Better", "OLS Better"), fill=c("grey","white"), xpd=NA, bty="n")
axis(2, labels = NA, tcl=-0.1)
axis(2, lwd = 0, line = -0.6, las=1)
box()
mtext("Relative Frequency", side = 2, line=-1.5, outer = TRUE, las=0, cex=1.2)
##===================


#======================================================
## Analysis 2: Sensitivity to Missing Data
#======================================================

##===================
## A) Simulating Replicate Datasets at Various Levels of Missing Data
##===================
# This analysis essentially repeats Analysis 1 but does so with varying amounts of missing data
# We want to simulate several levels of data missingness, but we want the final datasets in each case to be the same size
# This way we can't attribute any differences in model performance to differences in statistical power
# At each level of data missingness, the resulting dataset will contain 30 pairs
# Those 30 pairs will represent ~95%%, 75%, 50%, 25%, 5% of the overall dataset
# The corresponding number of species in each tree will then be:
# 9 species --> 36 pairs --> 30 pairs is 83.33% of all possible pairs
# 10 species --> 45 pairs --> 30 pairs is 66.66% of all possible pairs
# 12 species --> 66 pairs --> 30 pairs is 45.45% of all possible pairs
# 16 species --> 120 pairs --> 30 pairs is 25% of all possible pairs
# 35 spciees --> 595 pairs --> 30 pairs is ~5% of all possible pairs

# Overall approach:
# Simulate some number (Nsim) of datasets with the full tree and fit the models to them
# For each dataset, randomly remove the necessary proportion of pairs and re-fit the models
# Calculate the percentage difference in mean slope values
# Note that this approach requires a substantial amount of computing resources (especially at a high number of pairs)

## The procedure is then: 
library(phytools)
library(MASS)
library(phylopairs)
Nsim=5 # for paper's analysis we used Nsim=500
ntaxa=c(9, 10, 12, 16, 35)
# Generate trees
trees = lapply(ntaxa, function(x) pbtree(n=x, nsim=Nsim))
# Generate a set of BM traits for each replicate tree
X=lapply(trees, function(x) lapply(x, fastBM))
# For each replicate trait dataset, find the pairwise combinations of pairs, their traits, and the square difference in their traits
xps = lapply(trees, function(x) lapply(x, function(y) data.frame(t(combn(y$tip.label,2)))))
Xs = lapply(X, function(x) lapply(x, function(y) t(combn(y, 2))))
ypairs = lapply(Xs, function(y) t(sapply(y, function(x) (x[,1]-x[,2])^2)))
# Calculate the sp.pair covariance matrix for pglmm and the two four random effect matrices for the Castillo (2017) method
cov.y=re.mats=list()
for(i in 1:length(ntaxa)) {
  cov.y[[i]]=re.mats[[i]]=list()
  for(j in 1:Nsim) {
    cov.y[[i]][[j]] = taxapair.vcv(sp.pairs=xps[[i]][[j]], tree=trees[[i]][[j]], spnames=TRUE) # this line is v. slow
    re.mats[[i]][[j]] = twoterm.plmm.mats(sp.pairs=xps[[i]][[j]], tree=trees[[i]][[j]])
  }
}
# Verify that each covariance matrix is valid
for(i in 1:length(ntaxa)) {
  print(all(sapply(re.mats[[i]], function(x) sapply(x[3:4], function(y) all(covmat.check(y))))))
  print(all(sapply(cov.y[[i]], function(x) all(covmat.check(x)))))
}
# Generate a predictor variable for each replicate dataset
pred = lapply(ypairs, function(x) t(replicate(Nsim, expr=rnorm(ncol(x)))))
# Put each set of predictor variables into a design matrix and join these matrices in a list
xmats = lapply(pred, function(x) apply(x, 1, function(y) cbind(rep(1, length(y)), y), simplify=F))
# Define an intercept and slope for the relationship between response and predictor
b0=1
b1=0.8
# Calculate predicted val of response variable based on predictor of each rep
resp.x = lapply(pred, function(x) b0 + b1*x)
# Calculate and add structured random effects and residuals to response
# First, generate, for each rep, random effects that are scaled versions of Y (a*Y, where Y=(xi-xj)^2)
sig2.rand = 0.5
u = lapply(ypairs, function(x) x*sig2.rand)
# Second, generate, for each rep, actual random error (epsilom)
sig2.eps=0.1
eps= lapply(ypairs, function(x) mvrnorm(n=Nsim, mu=rep(0, ncol(x)), Sigma=diag(ncol(x))*sig2.eps))
# Compile these effects together to get an 'observed' response variable
resp=list()
for(i in 1:length(ntaxa)) resp[[i]] = resp.x[[i]] + u[[i]] + eps[[i]]
# Visualize the relationship between response and predictor
plot(pred[[5]][5,], resp[[5]][5,])
# Condense these datasets to just the node-averaged values
node.av=node.av.wtd=list()
for(i in 1:length(ntaxa)) {
	node.av[[i]]=node.av.wtd[[i]]=list()
	for(j in 1:Nsim) {
	  set = cbind(xps[[i]][[j]][,1:2], predic=pred[[i]][j,], respon=resp[[i]][j,])
	  pred.nodes=node.averager(tree=trees[[i]][[j]], taxacolumns=c("X1","X2"), dataset=set, varb="predic")
	  resp.nodes=node.averager(tree=trees[[i]][[j]], taxacolumns=c("X1","X2"), dataset=set, varb="respon")
	  pred.nodes.wtd=node.averager(tree=trees[[i]][[j]], taxacolumns=c("X1","X2"),dataset=set, varb="predic", weighted=T)
	  resp.nodes.wtd=node.averager(tree=trees[[i]][[j]], taxacolumns=c("X1","X2"),dataset=set, varb="respon", weighted=T)
	  node.av.mat=cbind(rep(1, length(pred.nodes)),pred.nodes)
	  node.av.wtd.mat=cbind(rep(1, length(pred.nodes.wtd)),pred.nodes.wtd)
	  node.av[[i]][[j]]=list(node.av.mat, resp.nodes)
	  node.av.wtd[[i]][[j]]=list(node.av.wtd.mat,resp.nodes.wtd)
    }
}
# Save the predictor variable design matrices, response variables, phylogenetic covariance matrices, and Z matrices
save(trees, xps, ypairs, xmats, resp, cov.y, re.mats, node.av, node.av.wtd, file="data.to.fit.missingtest.full.rdata")
# save(xmats, resp, cov.y, file="data.to.fit.missingtest.full.zeroslope.rdata") # if testing fits to data with no relationship

## Now, for each replicate, get rid of some number of pairs and re-calculate the corresponding matrices
# Draw 30 random indices for the pairs to keep for each replicate dataset
inds = lapply(xps, function(x) t(replicate(Nsim, sort(sample(nrow(x[[1]]), 30)))))
# Cut each dataset down to just these pairs (will have to recalculate the Z matrices for the Castillo (2017) method and the node averages in the next step)
# This is ugly code, but so be it:
xmats.cut=resp.cut=covy.cut=xps.cut=remats.cut=node.av.cut=node.av.wtd.cut=list()
for(i in 1:length(inds)) {
	xmats.cut[[i]]=covy.cut[[i]]=xps.cut[[i]]=remats.cut[[i]]=node.av.cut[[i]]=node.av.wtd.cut[[i]]=list()
	resp.cut[[i]] = matrix(NA,nrow(inds[[i]]),ncol(inds[[i]]))
	for(j in 1:nrow(inds[[i]])) {
		ind=inds[[i]][j,]
		xmats.cut[[i]][[j]] = xmats[[i]][[j]][ind,]
		covy.cut[[i]][[j]] = cov.y[[i]][[j]][ind,ind]
		resp.cut[[i]][j,] = resp[[i]][j,ind]
		xps.cut[[i]][[j]] = xps[[i]][[j]][ind,]
        remats.cut[[i]][[j]] = twoterm.plmm.mats(sp.pairs=xps.cut[[i]][[j]], tree=trees[[i]][[j]])
        set = cbind(xps.cut[[i]][[j]][,1:2], predic=xmats.cut[[i]][[j]][,2], respon=resp.cut[[i]][j,])
        pred.nodes=node.averager(tree=trees[[i]][[j]], taxacolumns=c("X1","X2"), dataset=set, varb="predic")
        resp.nodes=node.averager(tree=trees[[i]][[j]], taxacolumns=c("X1","X2"), dataset=set, varb="respon")
        pred.nodes.wtd=node.averager(tree=trees[[i]][[j]], taxacolumns=c("X1","X2"),dataset=set, varb="predic", weighted=T)
	    resp.nodes.wtd=node.averager(tree=trees[[i]][[j]], taxacolumns=c("X1","X2"),dataset=set, varb="respon", weighted=T)
        node.av.mat=cbind(rep(1, length(pred.nodes)),pred.nodes)
        node.av.wtd.mat=cbind(rep(1, length(pred.nodes.wtd)),pred.nodes.wtd)
        node.av.cut[[i]][[j]]=list(unname(node.av.mat), resp.nodes)
	    node.av.wtd.cut[[i]][[j]]=list(unname(node.av.wtd.mat),resp.nodes.wtd)
	}
}
save(trees, xps.cut, xmats.cut, resp.cut, covy.cut, remats.cut, node.av.cut, node.av.wtd.cut, 
	node.av.cut.unpruned, node.av.wtd.cut.unpruned, file="data.to.fit.missingtest.clipped.rdata")
##===================

##===================
## B) Fitting Models to the Replicate Full-Sized and Clipped Datasets
##===================
library(parallel)
library(phylopairs)
## Models should be fit to each replicate full-sized datasets and to replicate at each level of data missingness
# The underlying approach to model fitting is identical to that shown in Analysis 1A. 
# How that approach will be scaled-up for the full number of replicates and missingness treatments will depend on the compute cluster being used
# Here we show how we did this in the case of the OLS model and using a laptop with 4 cores 
# Other models take considerably longer to fit and necessitate the use of a cluster
# For code on how this was exactly done for the 'longleaf' cluster at UNC Chapel Hill in 2023/2024,

## i) Missing Data Fitting
load("data.to.fit.missingtest.clipped.rdata"))
# Organize the relevant data vectors into a list of lists
veclist.ols = list()
for(i in 1:5) {
  veclist.ols[[i]] = list()
  for(j in 1:Nsim) {
   veclist.ols[[i]][[j]] = list(xmats.cut[[i]][[j]], resp.cut[[i]][j,])
  }
}
# missingness level 1
outs.ols.missing.clipped.1 = mclapply(veclist.ols[[1]], function(x) linreg.stan(des=x[[1]], y=x[[2]]), mc.cores=4)
outs.node.missing.clipped.1 = mclapply(node.av.cut[[1]], function(x) linreg.stan(des=x[[1]][!is.na(x[[1]][,2]),], y=x[[2]][!is.na(x[[1]][,2])]), mc.cores=4)
outs.node.wtd.missing.clipped.1 = mclapply(node.av.wtd.cut[[1]], function(x) linreg.stan(des=x[[1]][!is.na(x[[1]][,2]),], y=x[[2]][!is.na(x[[1]][,2])]), mc.cores=4)
# missingness level 2
outs.ols.missing.clipped.2 = mclapply(veclist.ols[[2]], function(x) linreg.stan(des=x[[1]], y=x[[2]]), mc.cores=4)
outs.node.missing.clipped.2 = mclapply(node.av.cut[[2]], function(x) linreg.stan(des=x[[1]][!is.na(x[[1]][,2]),], y=x[[2]][!is.na(x[[1]][,2])]), mc.cores=4)
outs.node.wtd.missing.clipped.2 = mclapply(node.av.wtd.cut[[2]], function(x) linreg.stan(des=x[[1]][!is.na(x[[1]][,2]),], y=x[[2]][!is.na(x[[1]][,2])]), mc.cores=4)
# missingness level 3
outs.ols.missing.clipped.3 = mclapply(veclist.ols[[3]], function(x) linreg.stan(des=x[[1]], y=x[[2]]), mc.cores=4)
outs.node.missing.clipped.3 = mclapply(node.av.cut[[3]], function(x) linreg.stan(des=x[[1]][!is.na(x[[1]][,2]),], y=x[[2]][!is.na(x[[1]][,2])]), mc.cores=4)
outs.node.wtd.missing.clipped.3 = mclapply(node.av.wtd.cut[[3]], function(x) linreg.stan(des=x[[1]][!is.na(x[[1]][,2]),], y=x[[2]][!is.na(x[[1]][,2])]), mc.cores=4)
# missingness level 4
outs.ols.missing.clipped.4 = mclapply(veclist.ols[[4]], function(x) linreg.stan(des=x[[1]], y=x[[2]]), mc.cores=4)
outs.node.missing.clipped.4 = mclapply(node.av.cut[[4]], function(x) linreg.stan(des=x[[1]][!is.na(x[[1]][,2]),], y=x[[2]][!is.na(x[[1]][,2])]), mc.cores=4)
outs.node.wtd.missing.clipped.4 = mclapply(node.av.wtd.cut[[4]], function(x) linreg.stan(des=x[[1]][!is.na(x[[1]][,2]),], y=x[[2]][!is.na(x[[1]][,2])]), mc.cores=4)
# missingness level 5
outs.ols.missing.clipped.5 = mclapply(veclist.ols[[5]], function(x) linreg.stan(des=x[[1]], y=x[[2]]), mc.cores=4)
outs.node.missing.clipped.5 = mclapply(node.av.cut[[5]], function(x) linreg.stan(des=x[[1]][!is.na(x[[1]][,2]),], y=x[[2]][!is.na(x[[1]][,2])]), mc.cores=4)
outs.node.wtd.missing.clipped.5 = mclapply(node.av.wtd.cut[[5]], function(x) linreg.stan(des=x[[1]][!is.na(x[[1]][,2]),], y=x[[2]][!is.na(x[[1]][,2])]), mc.cores=4)

# Full-Sized Dataset Fitting
load("data.to.fit.missingtest.full.rdata")
veclist.ols.full = list()
for(i in 1:5) {
  veclist.ols.full[[i]] = list()
  for(j in 1:Nsim) {
   veclist.ols.full[[i]][[j]] = list(xmats[[i]][[j]], resp[[i]][j,])
  }
}
# level 1
outs.ols.missing.full.1 = mclapply(veclist.ols.full[[1]], function(x) linreg.stan(des=x[[1]], y=x[[2]]), mc.cores=4)
outs.node.missing.full.1 = mclapply(node.av[[1]], function(x) linreg.stan(des=x[[1]], y=x[[2]]), mc.cores=4)
outs.node.wtd.missing.full.1 = mclapply(node.av.wtd[[1]], function(x) linreg.stan(des=x[[1]], y=x[[2]]), mc.cores=4)
# level 2
outs.ols.missing.full.2 = mclapply(veclist.ols.full[[2]], function(x) linreg.stan(des=x[[1]], y=x[[2]]), mc.cores=4)
outs.node.missing.full.2 = mclapply(node.av[[2]], function(x) linreg.stan(des=x[[1]], y=x[[2]]), mc.cores=4)
outs.node.wtd.missing.full.2 = mclapply(node.av.wtd[[2]], function(x) linreg.stan(des=x[[1]], y=x[[2]]), mc.cores=4)
# level 3
outs.ols.missing.full.3 = mclapply(veclist.ols.full[[3]], function(x) linreg.stan(des=x[[1]], y=x[[2]]), mc.cores=4)
outs.node.missing.full.3 = mclapply(node.av[[3]], function(x) linreg.stan(des=x[[1]], y=x[[2]]), mc.cores=4)
outs.node.wtd.missing.full.3 = mclapply(node.av.wtd[[3]], function(x) linreg.stan(des=x[[1]], y=x[[2]]), mc.cores=4)
# level 4
outs.ols.missing.full.4 = mclapply(veclist.ols.full[[4]], function(x) linreg.stan(des=x[[1]], y=x[[2]]), mc.cores=4)
outs.node.missing.full.4 = mclapply(node.av[[4]], function(x) linreg.stan(des=x[[1]], y=x[[2]]), mc.cores=4)
outs.node.wtd.missing.full.4 = mclapply(node.av.wtd[[4]], function(x) linreg.stan(des=x[[1]], y=x[[2]]), mc.cores=4)
# level 5
outs.ols.missing.full.5 = mclapply(veclist.ols.full[[5]], function(x) linreg.stan(des=x[[1]], y=x[[2]]), mc.cores=4)
outs.node.missing.full.5 = mclapply(node.av[[5]], function(x) linreg.stan(des=x[[1]], y=x[[2]]), mc.cores=4)
outs.node.wtd.missing.full.5 = mclapply(node.av.wtd[[5]], function(x) linreg.stan(des=x[[1]], y=x[[2]]), mc.cores=4)
##===================

##===================
## C) Summarizing Results
##===================
# Here we show the results from the OLS model fits whose code is shown above. For the paper's analysis, the exact same procedure was used for all other model fits. 
reldiff = function(clipped, full, truth=0.8) {
	res = 100*(clipped[[1]][2,"mean"] - full[[1]][2,"mean"])/truth
	return(res)
}
ols.perc.diffs1=ols.perc.diffs2=ols.perc.diffs3=ols.perc.diffs4=ols.perc.diffs5=rep(NA, Nsim)
for(i in 1:Nsim) {
  ols.perc.diffs1[i] = reldiff(clipped=outs.ols.missing.clipped.1[[i]], full=outs.ols.missing.full.1[[i]])
  ols.perc.diffs2[i] = reldiff(clipped=outs.ols.missing.clipped.2[[i]], full=outs.ols.missing.full.2[[i]])
  ols.perc.diffs3[i] = reldiff(clipped=outs.ols.missing.clipped.3[[i]], full=outs.ols.missing.full.3[[i]])
  ols.perc.diffs4[i] = reldiff(clipped=outs.ols.missing.clipped.4[[i]], full=outs.ols.missing.full.4[[i]])
  ols.perc.diffs5[i] = reldiff(clipped=outs.ols.missing.clipped.5[[i]], full=outs.ols.missing.full.5[[i]])
}
boxplot(ols.perc.diffs1, ols.perc.diffs2, ols.perc.diffs3, ols.perc.diffs4, ols.perc.diffs5, ylim=c(-300,200), 
	names=c("16.67", "33.33", "54.54", "75.00", "94.06"))
abline(h=0, col="red")
mtext("Difference in Slope Estimate Relative to True Slope", side = 2, line = -1.5, outer=T, cex=1.2)
mtext("Percentage of Possible Pairs Missing", side = 1, line = -2.5, outer=T, cex=1.2)
##===================

##===================
## D) Tests of Bias Induced by Node-Averaging Procedures
##===================
# Here we calculate node averages for the same tree based on various amounts of missing data
# We do this for both unweighted and weighted node-averaging procedures
# Load packages
library(phytools)
library(parallel)
library(phylopairs)

# Define function for calculating the age of each node in the tree
node.age.getter = function(tree, dataset, taxacolumns=c("sp1.matched","sp2.matched")) {
# ensure the dataset has the right column names
  if(all(c(taxacolumns) %in% colnames(dataset))==F) {
    stop("one or more required columns is missing in the dataset, or there is an incorrect column name")
  }
  sp1=dataset[,taxacolumns[1]]
  sp2=dataset[,taxacolumns[2]]
  # Prune the tree to have just the taxa that are in the dataset (required due to extreme size and number of internal nodes of some trees)
  tree = drop.tip(phy=tree, tip=tree$tip.label[which(tree$tip.label %in% c(sp1,sp2)==F)])
  # calculate ages for the nodes in the tree
  nodes = round(max(node.depth.edgelength(tree)) - node.depth.edgelength(tree),4)
  res = nodes[nodes>0]
  return(res)
}

# Define the number of taxa to generate in the tree, the number of replicate datasets to create, and the number of cores to use when running code in parallel
Ntaxa=20
Nrep=1000
ncores=8

# Simulate a tree (we need to do this with one tree to keep the node ages the same) 
tree = pbtree(b=1, d=0, n=Ntaxa, type="continuous")

# Generate a table with all the possible pairwise crosses for the taxa in the simulated tree (table will have choose(Ntaxa,2) rows)
pairwise.sp=data.frame(t(combn(tree$tip.label,2)))
colnames(pairwise.sp)=c("sp1","sp2")

# Find the "age" of each node in the tree
node.ages = node.age.getter(tree=tree, dataset=pairwise.sp, taxacolumns=c("sp1","sp2"))

# Check that the node ages look correct when aligned to the tree plotted with a scale
plot(tree)
axisPhylo()
nodelabels(text = round(node.ages, 2), frame = "n")

# Make Nrep new tables of pairwise.sp plus a column with random positive values; these represent some hypothetical pairwise-defined trait (e.g. RI, geographic overlap, midpoint latitude)
pairwise.sp=rep(list(pairwise.sp),Nrep)
for(i in 1:Nrep) pairwise.sp[[i]]$trait = abs(rnorm(nrow(pairwise.sp[[1]]), mean=5))

# Apply Fitz and Turelli's correction to get "weighted" average values at each node in the tree and Coyne and Orr's (1989) correction to get a "unweighted" average value
trait.corrected = mclapply(pairwise.sp, function(x) node.averager(tree=tree, dataset=x, taxacolumns=c("sp1","sp2"), varb="trait", weighted=T), mc.cores=ncores)
trait.corrected.unwtd = mclapply(pairwise.sp, function(x) node.averager(tree=tree, dataset=x, taxacolumns=c("sp1","sp2"), varb="trait"), mc.cores=ncores)

# Sanity check: plot some corrected values
ind=10
plot(x=node.ages, y=trait.corrected[[ind]], xlab="node age", ylab="corrected val") 
plot(x=node.ages, y=trait.corrected.unwtd[[ind]], xlab="node age", ylab="corrected val") 
# These should show the homogenizing effect of the correction (i.e. values at deeper nodes should be less extreme than those at shallow nodes due to averaging)

# Define the levels of missing data 
# note: depending on the size of the tree and dataset, these percentages won't always generate whole numbers, so make sure to round up in the next step
levs = c(0.05, 0.15, 0.25, 0.5, 0.75, 0.9) 

# Generate Nrep datasets for each of the different levels of missing data
# For each replicate dataset, at each level of missing data, remove random rows of pairwise data
datasets=list()
for(i in 1:length(levs)) {
  datasets[[i]]=list()
  for(j in 1:Nrep) {
    inds = sample(x=1:nrow(pairwise.sp[[1]]), ceiling(levs[i]*nrow(pairwise.sp[[1]])))
    res=pairwise.sp[[j]]
    res[inds,"trait"]=NA
    datasets[[i]][[j]] = res
  }
}

# Double-check that the number of pairwise rows WITH data is the same in each replicate dataset (with the same level of missing data)
sapply(datasets, function(x) sapply(x, function(y) nrow(y[is.na(y$trait)==F,]))) # these are all the same
# What proportions of the original dataset to these values add up to? 
props = round(100*sapply(datasets, function(y) nrow(y[[1]][is.na(y[[1]]$trait)==F,]))/nrow(pairwise.sp[[1]]))
props

# Apply weighted and unweighted node averaging to each dataset that has missing values (can take a minute or two if running in parallel, several minutes if in serial)
corr.res=corr.res.unwtd=list()
for(i in 1:length(levs)) {
    corr.res[[i]] = mclapply(datasets[[i]], function(x) node.averager(tree=tree, dataset=x, taxacolumns=c("sp1","sp2"), varb="trait", weighted=T),mc.cores=ncores)
    corr.res.unwtd[[i]] = mclapply(datasets[[i]], function(x) node.averager(tree=tree, dataset=x, taxacolumns=c("sp1","sp2"), varb="trait"),mc.cores=ncores)
}

# Calculate and summarize the differences in the values calculated at each node when data are missing versus the full dataset for both methods
perc.missing = data.frame(matrix(NA, 4, length(node.ages)))
rownames(perc.missing) = c("mean","median","iqlow","iqhigh")
perc.missing=perc.missing.unwtd=rep(list(perc.missing), 6) 
for(i in 1:length(perc.missing)) {
  res = res.unwtd = matrix(NA, nrow=length(node.ages), ncol=Nrep)
  for(j in 1:Nrep) {
    res[,j] = 100*(corr.res[[i]][[j]] - trait.corrected[[j]])/trait.corrected[[j]]
    res.unwtd[,j] = 100*(corr.res.unwtd[[i]][[j]] - trait.corrected.unwtd[[j]])/trait.corrected.unwtd[[j]]
  }
  perc.missing[[i]] = apply(res, 1, function(x) summary(x)[c(4,3,2,5)])
  perc.missing.unwtd[[i]] = apply(res.unwtd, 1, function(x) summary(x)[c(4,3,2,5)])
}

# Plot percentage differece in estimates from different datasets 
# Note that this might not look exactly like that shown in the paper due to stochasticity, but overall pattern should be the same
par(mfrow=c(1,2))
par(mar=c(8, 3, 8, 0), xpd=TRUE)
cols=c("blue", "green", "orange", "purple", "brown")
plot(x=node.ages, y=perc.missing[[1]][2,], ylim=c(-100, 10), pch=19, yaxt='n', xaxt='n', xlab='')
axis(2, labels = NA)
axis(2, lwd = 0, line = -0.4, las=0)
axis(1, labels = NA)
axis(1, lwd = 0, line = -0.4, las=0)
lines(node.ages[order(node.ages)], perc.missing[[1]][2,][order(node.ages)])
abline(h=0, lty="dotted", col="gray60", xpd=F)
for(j in 2:length(levs)) {
  points(x=node.ages, y=perc.missing[[j]][2,], col=cols[j-1], pch=19)
  lines(node.ages[order(node.ages)], perc.missing[[j]][2,][order(node.ages)], col=cols[j-1])
}
mtext("Percent Difference in Node Average", side = 2, line = -1, outer = TRUE, las=0, cex=1.2)
par(mar=c(8, 0.8, 8, 2.2), xpd=TRUE)
plot(x=node.ages, y=perc.missing.unwtd[[length(levs)]][2,], ylim=c(-100, 10), 
  pch=19, yaxt='n', xaxt='n', xlab='', col=cols[length(levs)-1])
axis(2, labels = NA)
axis(1, labels = NA)
axis(1, lwd = 0, line = -0.4, las=0)
lines(node.ages[order(node.ages)], perc.missing.unwtd[[length(levs)]][2,][order(node.ages)])
abline(h=0, lty="dotted", col="gray60", xpd=F)
mtext("Node Depth (Time from Present)", side = 1, line = -6, outer = TRUE, las=0, cex=1.2)
legend("bottomright", legend=100-props, col=c("black", cols), pch=19, title="Data Missing (%)")

## Now add in the empirical data from Price Bouvier (2002)
## See the file anderson.etal.datawrangling.R for more information on these data
# Load tree and dataset
birdtree = read.tree("birds_pulidoSC2016_MaxCladeCredTree.txt")
birdtree 
pb.in.tree=read.csv("pb.in.tree.csv", stringsAsFactor=F)
# Calculate node ages and node-averaged RI values
node.ages = node.age.getter(tree=birdtree, dataset=pb.in.tree, taxacolumns=c("sp1","sp2"))
wtd.ri = node.averager(tree=birdtree, dataset=pb.in.tree, taxacolumns=c("sp1","sp2"), weighted=T, varb="fertility")
unwtd.ri = node.averager(tree=birdtree, dataset=pb.in.tree, taxacolumns=c("sp1","sp2"), varb="fertility")
# First get a sense of the relationship between node age and the weighted node-averaged value of strength of RI
plot(node.ages, wtd.ri, pch=16)
# Soom into relevant x-axis values
plot(node.ages[!is.na(wtd.ri)], wtd.ri[!is.na(wtd.ri)], pch=16)
# Add the OLS regression line to the graph
abline(lm(wtd.ri[!is.na(wtd.ri)]~node.ages[!is.na(wtd.ri)]), col="red", lwd=3)
# Check the OLS statistics
summary(lm(wtd.ri[!is.na(wtd.ri)]~node.ages[!is.na(wtd.ri)]))
# This should reveal a marginally insignificant negative relationship between node age and strength of RI
# Now compare this with the OLS statistics for the relationship between the raw measurements for strength of RI and node age 
summary(lm(fertility~pair.age, data=pb.in.tree)) 
# This should reveal a highly significant pos. relationship in the raw data
# Plot a comparison between raw data and the two forms of node-averaged data
dev.new()
par(mfrow=c(1,3))
plot(pb.in.tree$pair.age, pb.in.tree$fertility, pch=16, yaxt='n', xlab='', ylab='', xaxt='n')
axis(2, labels = NA)
axis(2, lwd = 0, line = -0.4, las=0)
axis(1, labels = NA)
axis(1, lwd = 0, line = -0.4, las=0)
mtext("Hybrid Fertility", side=2, line=2)
mtext("Age of Species in Cross (Ma)", side=1, line=2)
abline(lm(pb.in.tree$fertility~pb.in.tree$pair.age), col="red", lwd=3)
plot(node.ages[!is.na(wtd.ri)], wtd.ri[!is.na(wtd.ri)], pch=16, yaxt='n', ylab='', xaxt='n', xlab='')
axis(2, labels = NA)
axis(2, lwd = 0, line = -0.4, las=0)
axis(1, labels = NA)
axis(1, lwd = 0, line = -0.4, las=0)
mtext("Weighted Average Hybrid Fertility", side=2, line=2)
mtext("Node Depth (Ma)", side=1, line=2)
abline(lm(wtd.ri[!is.na(wtd.ri)]~node.ages[!is.na(wtd.ri)]), col="red", lwd=3)
plot(node.ages[!is.na(unwtd.ri)], unwtd.ri[!is.na(unwtd.ri)], pch=16, yaxt='n', ylab='', xaxt='n', xlab='')
axis(2, labels = NA)
axis(2, lwd = 0, line = -0.4, las=0)
axis(1, labels = NA)
axis(1, lwd = 0, line = -0.4, las=0)
mtext("Node Depth (Ma)", side=1, line=2)
mtext("Uneighted Average Hybrid Fertility", side=2, line=2)
abline(lm(unwtd.ri[!is.na(unwtd.ri)]~node.ages[!is.na(unwtd.ri)]), col="red", lwd=3)
##===================


#======================================================
## Analysis 3: Parameter Estimation, Bounded Response
#======================================================

##===================
## A) Simulating Replicate Datasets of Bounded Pairwise-Defined Traits
##===================
# The procedure for simulating these data is identical to the procedure from Analysis 1 up until line 65
# That is: replicate trees, pairs, BM-evolving traits, covariance matrices, Z matrices, predictor variables, and predictor design matrices are the same 
# The only differences are:
# 1. now the response is bounded instead of unbounded, and 
# 2. the linear relationship is on the scale of a link function. 

# Assuming the objects created up until line 65 of this document are in the global environment, the remaining simulation procedure is as follows: 
# Define parameters: an intercept and slope for the relationship between response and predictor on the scale of the linear predictor, and a sig2.rand scaling parameter
b0=0
b1=2 
sig2.rand=1
## NOTE: for the analysis in the paper, we used all combinations of the following parameters: 
## b1 = 0, 0.8, 2
## sig2.rand = 0.01, 0.10, 1.00
# Calculate predicted val of response on the scale of the linear predictor based on the predictor variable of each rep
resp.x = b0 + b1*pred
# Calculate and add structured random effects to response
u = t(sapply(cov.y, function(x) MASS::mvrnorm(n=1, mu=rep(0, nrow(x)), Sigma=x*sig2.rand)))
## NOTE: in the paper's analysis of this model, we also generated beta-distributed response variables in which there was no covariance among observations. 
## To do so, all code to this point is the same, except the covariance matrix used in the above line of code becomes an identity matrix
## The code above becomes: 
## u.uncorr = t(sapply(cov.y, function(x) MASS::mvrnorm(n=1, mu=rep(0, nrow(x)), Sigma=diag(nrow(x))*sig2.rand)))
## We code it like this to make the two lines of code directly comparable, but the same can be coded in many other, faster ways
# Calculate what the link function of the predicted y would be, given the predicted val of the response and the random effects, and assuming a logit link function
logit.mu = resp.x + u
# Calculate what the actual predicted y is from the inverse of the link function 
mu = exp(logit.mu)/(1 + exp(logit.mu)) 
# Define precision parameter phi
phi = 5
# Convert mu and phi to alpha and beta, which are the beta-distribution parameters recognized in R
alpha.par = mu*phi
beta.par = (1 - mu)*phi
# Simulate the response variable from beta distribution
resp = matrix(rbeta(length(mu), shape1 = alpha.par, shape2 = beta.par), ncol=ncol(mu))
# Visualize the relationships to make sure they don't look wonky
ind=1
plot(pred[ind,], resp.x[ind,]) # should be linear and deterministic, y unbounded
plot(pred[ind,], logit.mu[ind,]) # should be linear with variation, y unbounded
plot(pred[ind,], resp[ind,]) # should be non-linear at tails, y bounded

# Now, just as in previous analyses, we use node averaging functions to condense these datasets to just the node-averaged values
node.av=node.av.wtd=list()
for(i in 1:Nsim) {
	set = cbind(xps[[i]][,1:2], pred[i,], resp[i,])
	pred.nodes=node.averager(tree=trees[[i]], taxacolumns=c("sp1","sp2"), dataset=set, varb="pred[i, ]")
	resp.nodes=node.averager(tree=trees[[i]], taxacolumns=c("sp1","sp2"), dataset=set, varb="resp[i, ]")
	pred.nodes.wtd=node.averager(tree=trees[[i]], taxacolumns=c("sp1","sp2"),dataset=set, varb="pred[i, ]", weighted=T)
	resp.nodes.wtd=node.averager(tree=trees[[i]], taxacolumns=c("sp1","sp2"),dataset=set, varb="resp[i, ]", weighted=T)
	node.av.mat=cbind(rep(1, length(pred.nodes)),pred.nodes)
	node.av.wtd.mat=cbind(rep(1, length(pred.nodes.wtd)),pred.nodes.wtd)
	node.av[[i]]=list(node.av.mat, resp.nodes)
	node.av.wtd[[i]]=list(node.av.wtd.mat,resp.nodes.wtd)
}

# Save the predictor variable design matrices, response variables, phylogenetic covariance matrices, and Z matrices depending on parameter scheme:
save(trees, xmats, resp, cov.y, re.mats, xps, node.av, node.av.wtd, file="beta.reg.data.to.fit.sig1.slp2.rdata")
##===================

##===================
## B) Fitting Models to Simulated Datasets 
##===================
# Fitting beta regression mixed-effect models to all replicate simulated datasets will require the use of a compute cluster
# Computing clusters vary in terms of how replicate jobs are run, so the code below is for fitting a model to a single replicate dataset
# This code can be modified and incorporated into scripts for fitting models to multiple replicate datsets in parallel
# For model-performance testing, we set iter=6000 for each model fitting function (where iter is the number of sampling iterations in the underlying Bayesian models)
library(phylopairs)
# Load dataset
load("beta.reg.data.to.fit.sig1.slp2.rdata")
# Define which replicate to run 
num=1
# Define the number of iterations to run
its=1000
# Fit a beta regression mixed-effects model to the dataset
outs.beta.mm = betareg.stan(des=xmats[[num]], y=resp[num,], model="beta.mm", cov=cov.y[[ind]], iter=its)
# Fit a standard beta regression model (that assumes no lineage-pair covariance) to the same dataset
outs.beta.ols = betareg.stan(des=xmats[[num]], y=resp[num,], model="beta.ols", iter=its)
# Observe parameter estimates (slope estimate is coef2; recall true slope is 2 in this example)
outs.beta.mm[[1]]
outs.beta.ols[[1]]
##===================


#======================================================
## Analysis 4: Empirical Example, Bird Hybridization Data
#======================================================
## Empirical analysis of Price and Bouvier (2002)
# Goal here is to fit alternative models to the data and determine which model is the best fit (via loo and the other methods)
# See anderson.etal.datawrangling.R for details on this dataset
# Note that due to performance issues identified by performance tests, we will not be conducting node-averaging procedures here.
require(ape)
require(phylopairs)
require(rstan)
# load tree
birdtree = read.tree("birds_pulidoSC2016_MaxCladeCredTree.txt")
birdtree 
# load datset
pb.in.tree=read.csv("pb.in.tree.csv", stringsAsFactor=F)

# Get a visual sense of the relationship between node age and ri values (repeating step from Analysis 2D)
plot(pb.in.tree$pair.age, pb.in.tree$fertility, pch=16, xlab="age of spp. in cross", ylab="RI")

## Fitting the models
## OLS on full data
pb.ols = linreg.stan(des=pb.in.tree$pair.age, y=pb.in.tree$fertility)
## The new lineage-pair LMM on full data
# First calculate the lineage-pair covariance matrix, then fit the model
lpcov = taxapair.vcv(sp.pairs=pb.in.tree[,1:2], tree=birdtree, spnames=FALSE, dec=2)
covmat.check(lpcov)
pb.lplmm = linreg.stan(des=pb.in.tree$pair.age, y=pb.in.tree$fertility, model="pgls.mm", covmat=lpcov, iter=6000)
## Castillo's (2017) two-term mixed-model approach on full data
pb.ttlmm = twoterm.plmm.stan(des=pb.in.tree$pair.age, y=pb.in.tree$fertility, sp1s=pb.in.tree$sp1, sp2s=pb.in.tree$sp2, tree=birdtree, iter=6000)
## Beta regression on full data
pb.beta = betareg.stan(des=pb.in.tree$pair.age, y=pb.in.tree$fertility/5, iter=6000)
## Beta mixed-effects model on full data
pb.beta.mm = betareg.stan(des=pb.in.tree$pair.age, y=pb.in.tree$fertility/5, model="beta.mm", covmat=lpcov, iter=6000)

## Visually compare estimated slopes from the models
ols.allo[[1]]
lplmm.allo[[1]]
ttlmm.allo[[1]]
beta.allo[[1]]
beta.mm.allo[[1]]

# loo
loo::loo(pb.ols$ll)
loo::loo(pb.lplmm$ll)
loo::loo(pb.ttlmm$ll)
loo::loo(pb.beta$ll)
loo::loo(pb.beta.mm$ll)

# waic
loo::waic(pb.ols$ll)
loo::waic(pb.lplmm$ll)
loo::waic(pb.ttlmm$ll)
loo::waic(pb.beta$ll)
loo::waic(pb.beta.mm$ll)


#======================================================
## Analysis 5: Empirical Example, Drosophila
#======================================================
# Goal here is to fit alternative models to the data and determine which model is the best fit (via loo and the other methods)
# See anderson.etal.datawrangling.R for details on this dataset
# Due to performance issues identified by performance tests, we will not be conducting node-averaging procedures here.
require(ape)
require(phylopairs)
require(rstan)
# Load tree
load("dros.ultra.rdata")
dros.ultra
# Load datset
dat=read.csv(file="yukilevich.dros.intree.csv", stringsAsFactor=F)

## There are dozens of analyses one might conceivably carry out with these data.
# Here we will (rather arbitrarily) be testing for an effect of ecological differentiation on sexual isolation in allopatric species pairs
# The sexual isolation metric is bounded by 0 and 1.
# Stan models cannot accept NAs. We will remove them here first, then zero-in on the allopatric pairs.
dat.na = dat[!(is.na(dat$sexual.isolation)|is.na(dat$Ecological.Distance)),]
dat.na.allos = dat.na[dat.na[,6]==0,] 
nrow(dat.na.allos)# 62 allopatric pairs.
# Note that one of these has a negative value for sexual.isolation, which is impossible.
# For now, remove this row
dat.na.allos = dat.na.allos[dat.na.allos$sexual.isolation>0,]
# OLS model
ols.allo = linreg.stan(des=dat.na.allos$Ecological.Distance, y=dat.na.allos$sexual.isolation)
# Lineage-pair LMM model
allo.cov = taxapair.vcv(sp.pairs=dat.na.allos[,c("Sp.1","Sp2")], tree=dros.ultra, spnames=FALSE, dec=2)
lplmm.allo = linreg.stan(des=dat.na.allos$Ecological.Distance, 
	y=dat.na.allos$sexual.isolation, model="pgls.mm", cov=allo.cov, iter=10000) # takes a few minutes when using four cores
# Two-term mixed-model
ttlmm.allo = twoterm.plmm.stan(des=dat.na.allos$Ecological.Distance, y=dat.na.allos$sexual.isolation, 
	sp1s=dat.na.allos$Sp.1, sp2s=dat.na.allos$Sp2, tree=dros.ultra, iter=6000) 
# Beta regression
beta.allo = betareg.stan(des=dat.na.allos$Ecological.Distance, y=dat.na.allos$sexual.isolation, iter=6000)
# error returned: the value for nasuta and albomicans in the dataset is negative, which is impossible. For now,remove that value.
# Beta mixed-effects model
beta.mm.allo = betareg.stan(des=dat.na.allos$Ecological.Distance, y=dat.na.allos$sexual.isolation, model="beta.mm", covmat=allo.cov, iter=6000)

## Visually compare estimated slopes from the models
ols.allo[[1]]
lplmm.allo[[1]]
ttlmm.allo[[1]]
beta.allo[[1]]
beta.mm.allo[[1]]

## Model selection
# loo
loo::loo(ols.allo$ll)
loo::loo(lplmm.allo$ll)
loo::loo(ttlmm.allo$ll)
loo::loo(beta.allo$ll)
loo::loo(beta.mm.allo$ll)

# waic
loo::waic(ols.allo$ll)
loo::waic(lplmm.allo$ll)
loo::waic(ttlmm.allo$ll)
loo::waic(beta.allo$ll)
loo::waic(beta.mm.allo$ll)
##===================
```
