## Supplementary data and code for "The comparative analysis of lineage-pair traits": anderson.etal.datawrangling.R

```
#==========================================
# Dataset 1: Avian hybridization data from Price and Bouvier (2002), https://doi.org/10.1111/j.0014-3820.2002.tb00133.x
#==========================================
# This dataset has been published upon numerous times
# I obtained the dataset from Jenn Coughlan on 25 April 2022; Coughlan and Matute had used the dataset in their 2020 Phil Trans paper (https://doi.org/10.1098/rstb.2019.0533)
# It's size appears to vary depending on source. 
# The dataset as obtained contains info on 404 crosses involving 368 species
# In a review paper by Edwards et al. (2005), they say that 254 crosses were looked at by Price and Bouvier
# Price and Bouvier report the larger number in their paper
# The ages of the splits for these crosses will be out of date 
# Here we wil use the Pulido-Santacruz et al. (2019) tree to update these age estimates
# Some notes from the paper worth considering: 
# "Nonindependence should not produce biases in our estimation of times to accumulate levels of postzygotic isolation, but precludes formal statistical testing."
# "In this analysis we are concerned with averages across classes of hybrid fitness, making deviations from a clock less critical."
# "Our use of a gamma-corrected distance metric should allow rough extrapolation over longer time scales, but eventually saturation effects will impact the calibration and we expect absolute dates over long time scales to be underestimated"

## Wrangling
# The original excel sheet was a .xls file (bouvier_price.xls)
# I made a second sheet in the excel dataset that simplified some of headings and focused on just the key data
# I saved the simplified version of the dataset as a csv file, price_bouvier.csv, so that I could do the rest in R and thus permit greater reproducibility
# Both files are included in the materials here. 
# It was impossible, however, to fully automate the renaming
# Problem was that the name inputs were quite variable and needed some manual standardization
# Specifically, for some species in the "Species 1" column, the subspecies name was not spelled out but was only given in letter form
# I manually replaced those letters with the spelled out subspecies names
# There were also two cases (one species in each column) that had an extra space between the names, which I manually removed
# Finally, there was a star next to the fertility score for a handful of pairs.
# I'm not sure what this star means and can't find any reference to it.
# I removed the star manually so that the data wouldn't have to be removed (the star makes that line come up as a character vector, and the star is converted to NAs if you just switch it to numerical)
# Once this was complete, the rest of the name standardizations could be done with the following code

# Load the file (we were to this as the pb dataset, for Price Bouvier)
pb=read.csv("price_bouvier.csv")
head(pb)

## Clean up the Dataset
## First, we want species names to be rewritten in the format Genus_species
# In the dataset I received, the names are separated by spaces instead of underscores
# In many cases, the species 2 column has just the genus letter (and the species letter in the case of subspecies)
# To reformat, first get rid of the periods after the genus or subspecies names in the sp2 column
pb$sp2 = gsub(".", "", pb$sp2, fixed=T)

# Now replace the genus letters in sp2 column with the genus names from the sp1 column, and do the same for subspecies letters
# Join the names by an underscore
for(i in 1:nrow(pb)) {
	k = strsplit(pb$sp2[i], " ")[[1]]
	j = strsplit(pb$sp1[i], " ")[[1]]
	if(length(k)==2 & nchar(k[1])==1) {
	  pb$sp2[i] = paste(j[1], k[2], sep="_")
    }
    if(length(k)==3 & nchar(k[1])==1 & nchar(k[2])==1) {
      pb$sp2[i] = paste(j[1], j[2], k[3], sep="_")
    }
    if(length(k)==3 & nchar(k[1])==1 & nchar(k[2])!=1) {
      pb$sp2[i] = paste(j[1], k[2], k[3], sep="_")
    }
}

# Replace the spaces between the taxonomic names to underscores in the remaining columns
pb$sp1 = gsub(" ", "_", pb$sp1)
pb$sp2 = gsub(" ", "_", pb$sp2)

# Ensure there's no weird errors in the naming -- maximum length of names should be three (when subspecies are present) and minimum should be two
summary(sapply(pb$sp1, function(x) length(strsplit(x, "_")[[1]])))
summary(sapply(pb$sp2, function(x) length(strsplit(x, "_")[[1]]))) # there's a entry with one name only
which(sapply(pb$sp2, function(x) length(strsplit(x, "_")[[1]]))==1) #303
pb[303,] # the genus name was dropped from sp2 here
pb[303,2]="Lonchura_striata"

# Inspecting the two columns, there's still a little cleaning up to do
# There was evidently an extra space at the end of some names in the original spreadsheet, so some species names have overhanging underscores after the binomial
# Remove those with the following lines
pb$sp1 = gsub("_$","",pb$sp1)
pb$sp2 = gsub("_$","",pb$sp2)

# =================================
require(ape)
require(diverge)
require(parallel)
require(phylopairs)
## TREES
# Tree 1: Pulido-Santacruz and Weir 2016 Evolution (referred to as psc-jtw tree)
# this tree is the Jetz et al. (2012) tree plus the Fernariidae tree from Derryberry et al. 2011 Evolution
# A sub-analysis of this paper looked at taxonomic orders or suborders if they contained enough species with genetic data (see table 1)
# the tree and other supplementary materials downloaded from dryad at https://datadryad.org/stash/dataset/doi:10.5061/dryad.2v462
# contains the MCC tree and 1000 trees sampled from the posterior distribution

## Load the tree
birdtree = read.tree("birds_pulidoSC2016_MaxCladeCredTree.txt")
birdtree # 9982 tips
# extract the unique species names from the Price Bouvier dataset
sps = unique(c(pb$sp1, pb$sp2))
# 381 species names in this pb dataset; how many of these species names are in the tree? 
length(which(sps %in% birdtree$tip.label)) #231
# clip the dataset to contain just those pairs for which both species are in the tree
pb.in.tree = pb[which(apply(pb[,1:2], 1, function(x) all(x %in% birdtree$tip.label))),]
nrow(pb.in.tree) # 207 pairs remaining
# Find the age of the mrca for each pair
pair.ages = pair.age.in.tree(dataset=pb.in.tree, tree=birdtree, taxacolumns=c("sp1","sp2"))
# Summarize 
pb.in.tree = cbind(pb.in.tree, pair.age = pair.ages)
# save this analysis-formatted dataset
write.csv(pb.in.tree, file="pb.in.tree.csv", row.names=F)

#==========================================
# Dataset 2: Drosophila hybridization data from drosophila-speciation-patterns.com, and originally from the references therein
#==========================================
## Empirical analysis of Drosophila data
# The dataset here is a slightly revised version of the "Raw Data Table" availale for download from https://life2.bio.sunysb.edu/drosophila/datatables.html
# That file was entitled RAW_datafile_October_23_2023.xlsx
# Before converting that file to a .csv file, I manually removed title rows -- that is, rows containing things like group names and subgroup names (these rows do not include data)
# I saved the resulting file as yukilevich.dros.csv
library(ape)

# Many of the formative papers on Drosophila hybridization were conducted decades ago and have badly outdated divergence times. 
# Moreover, phylogenetic relationships were only known within small sets of subgroups.
# Today, there is much greater information on phylogenetic relationships among Drosophilids
# Our first goal is to update the phylogenetic relationships from the newest Drosophila phylogeny
# That phylogeny is from Kim et al. 2024, PLoS Biology https://journals.plos.org/plosbiology/article?id=10.1371/journal.pbio.3002697
# Load the new tree from Kim et al. paper
dros=read.tree("full_scaled_4d.tree")
# Load the dataset
dat=read.csv("yukilevich.dros.csv")
dim(dat) #610*132

## Prepping the Dataset
# Lots of the species names are divided into sub-species or geographic races -- these taxa will not be in the phylogenetic tree
# Our purposes here are to illustrate a use-case of the new models, so we can keep this simple and cut the dataset down to just rows that contain full species
# More in-depth analyses of certain hypotheses for Drosophila speciation should make a greater effor to reconcile species and tree such that more data are retained
k=strsplit(dat$Sp.1, "[ ]")
f=sapply(k, length)
k=strsplit(dat$Sp2, "[ ]")
g=sapply(k, length)
dat=dat[which(f==1&g==1),]
# One column uses the Sobel and Chen 2014 metric of sexual isolation
# This column has a cumbersome name in the dataset and uses "Chang" instead of Chen
# Renaming this; it will be the response variable here
colnames(dat)[13]="sexual.isolation"
# Make the numerical values numeric
dat[,13]=as.numeric(dat[,13])
dat[,6]=as.numeric(dat[,6])
dat$Ecological.Distance=as.numeric(dat$Ecological.Distance)
dat[,47]=as.numeric(dat[,47]) # mean lat sp. 1
dat[,48]=as.numeric(dat[,48]) # mean lat sp. 2
mean.lat=(abs(dat[,47])+abs(dat[,48]))/2
lat.diff=abs(dat[,47]-dat[,48])

## How many of the species in the dataset are in the tree? 
# The tree includes genus names, the dataset doesn't
# The genus names in the tree are shortened to one or a couple letters instead of full names (e.g. D for Drosophila)
spnames=strsplit(dros$tip.label, "[.]")
dros.genus = which(sapply(spnames, function(x) x[1]=="D"))
spnames=sapply(spnames[dros.genus], function(x) x[2])
m=which(unique(c(dat$Sp.1, dat$Sp2)) %in% spnames)
length(m)# 112 species in the dataset are also in the tree
# More importantly, how many pairs in the dataset have both species in the tree? 
intree = which(dat$Sp.1 %in% spnames & dat$Sp2 %in% spnames) 
length(intree)# 150 total pairs
# We'll clip the dataset to contain just these pairs
dat = dat[intree,]

## Prepping the Tree
# Going forward, the names in the dataset and in the tree will have to match, so I'll rename the species in the tree
dros$tip.label[dros.genus]=spnames
# Importantly, this tree, while containing the most comprehensive and up-to-date estimates of drosophilid relationships currently available, is not ultrametric
# We need to make this tree ultrametric, otherwise the lineage-pair covariance matrix won't be a valid covariance matrix
# (i.e. it won't be positive definite and it won't be diagonally dominant)
# Again, given the purposes of this analysis (demonstrating model fits), a faily quick-and-dirty approach is ok here
# We will make an ultrametric tree with branches in relative time (we don't have calibration points) by using the makeChronosCalib function of ape
# Using this function requires a bit of trial and error in terms of parameters: 
dros.relative = makeChronosCalib(dros, node = "root", age.min = 1, age.max = 1, interactive = FALSE, soft.bounds = FALSE) 
# We can test out different lambda values, which controls evolutionary rate variation among branches (lambda = 1 means constant rate, lambda = 0 means highly variabel rate)
dros1 = chronos(dros, model = "relaxed", lambda =0.1, calibration = dros.relative, control = chronos.control(maxit = 10000)) 
dros2 = chronos(dros, model = "relaxed", lambda =1, calibration = dros.relative) 
dros3 = chronos(dros, model = "relaxed", lambda =10, calibration = dros.relative) 
dros4 = chronos(dros, model = "relaxed", lambda =0, calibration = dros.relative) 
# The final value gives the best PHIIC and log-like
# Now recalculate the lineage-pair covariance matrix 
dros.ultra=dros4
plot(dros.ultra, cex=0.2)

## Save the files
# For downstream analyses, we'll use the condensed and renamed trait dataset and the relative time ultrametric tree
save(dros.ultra, file="dros.ultra.rdata")
write.csv(dat, file="yukilevich.dros.intree.csv")
```
