## Supplementary data and code for "The comparative analysis of lineage-pair traits": README.txt

This folder contains code and data files for conducting the analyses in Anderson et al. SUBMITTED.

Data files include raw data files as in the formats in which they were downloaded as well as the analysis-ready files that were created from the raw files. Code for converting raw data to analysis-ready formats is given in anderson.etal.datawrangling.R. 

The files that should appear in this folder are as follows. 

Downloaded Data Files:
1. full_scaled_4d.tree: Drosophila tree from Kim et al. 2024 PLoS Biology
2. birds_pulidoSC2016_MaxCladeCredTree.txt: All-birds phylogeny from Pulido-Santacruz and Weir 2016 Evolution. 
3. bouvoir_price.xls: Avian hybridization data from Price and Bouvier (2002)
4. price_bouvier.csv: Avian hybridization data from Price and Bouvier (2002) converted to .csv format
5. RAW_datafile_October_23_2023.xlsx: Fly hybridization data as downloaded from drosophila-speciation-patterns.com
6. yukilevich.dros.csv: Fly hybridization data converted into .csv format

Analysis-Ready Files: 
1. yukilevich.dros.intree.csv: Fly hybridization dataset for which subspecies and geographic races have been removed, among other processing details (see the data wrangling R file for details). 
2. dros.ultra.rdata: Ultra metric relative-time version of the Kim et al. (2024) Drosophila tree. 
3. pb.in.tree.csv: Price-Bouvier data trimmed to contain just the crosses for which both species are included in the Pulido-Santacruz et al. phylogenetic tree. 

Code files: 
anderson.etal.datawrangling.R: Code for converting downloaded data files to analysis-ready files
anderson.etal.analyses.R: Code for conducting model performance tests and re-analyses of empirical bird and drosophila datasets. 
